# Predictive representations can link model-based reinforcement learning to model-free mechanisms

**DOI:** 10.1101/083857

**Authors:** Evan M. Russek, Ida Momennejad, Matthew M. Botvinick, Samuel J. Gershman, Nathaniel D. Daw

## Abstract

Humans and animals are capable of evaluating actions by considering their long-run future rewards through a process described using model-based reinforcement learning (RL) algorithms. The mechanisms by which neural circuits perform the computations prescribed by model-based RL remain largely unknown; however, multiple lines of evidence suggest that neural circuits supporting model-based behavior are structurally homologous to and overlapping with those thought to carry out model-free temporal difference (TD) learning. Here, we lay out a family of approaches by which model-based computation may be built upon a core of TD learning. The foundation of this framework is the successor representation, a predictive state representation that, when combined with TD learning of value predictions, can produce a subset of the behaviors associated with model-based learning, while requiring less decision-time computation than dynamic programming. Using simulations, we delineate the precise behavioral capabilities enabled by evaluating actions using this approach, and compare them to those demonstrated by biological organisms. We then introduce two new algorithms that build upon the successor representation while progressively mitigating its limitations. Because this framework can account for the full range of observed putatively model-based behaviors while still utilizing a core TD framework, we suggest that it represents a neurally plausible family of mechanisms for model-based evaluation.

**Author Summary:** According to standard models, when confronted with a choice, animals and humans rely on two separate, distinct processes to come to a decision. One process deliberatively evaluates the consequences of each candidate action and is thought to underlie the ability to flexibly come up with novel plans. The other process gradually increases the propensity to perform behaviors that were previously successful and is thought to underlie automatically executed, habitual reflexes. Although computational principles and animal behavior support this dichotomy, at the neural level, there is little evidence supporting a clean segregation. For instance, although dopamine — famously implicated in drug addiction and Parkinson’s disease — currently only has a well-defined role in the automatic process, evidence suggests that it also plays a role in the deliberative process. In this work, we present a computational framework for resolving this mismatch. We show that the types of behaviors associated with either process could result from a common learning mechanism applied to different strategies for how populations of neurons could represent candidate actions. In addition to demonstrating that this account can produce the full range of flexible behavior observed in the empirical literature, we suggest experiments that could detect the various approaches within this framework.

## Introduction

A key question in both neuroscience and psychology is how the brain evaluates candidate actions in complex, sequential decision tasks. In principle, computing an action’s expected long-run cumulative future reward (or *value*) requires averaging rewards over the many future state trajectories that might follow the action. In practice, the exact computation of such expectations by dynamic programming or tree search methods may be prohibitively expensive, and it is widely believed that the brain simplifies the computations occurring at decision time, in part by relying on “cached” (pre-computed) long-run value estimates [1].

Such caching of values is the hallmark of prominent temporal difference (TD) learning theories, according to which prediction errors reported by phasic dopamine responses update these cached variables [2–4]. This, in turn, provides a neuro-computational account of inflexible stimulus-response habits, due to the fact that TD learning cannot update cached values in response to distal changes in reward (e.g., following reward devaluation). The computationally cheap but inflexible “model-free” nature of TD learning, which relies only on trial-and-error interactions with the environment, contrasts with the flexible but computationally expensive “model-based” nature of dynamic programming and tree search methods, which rely on an explicit internal model of the environment. Due to the complementary properties of model-free and model-based value computation, it has been suggested that the brain makes use of both in the form of parallel RL systems that compete for control of behavior [1].

Although flexible choice behavior seems to demonstrate that humans and animals may use model-based evaluation in some circumstances, very little is known about how this is actually implemented in the brain, or indeed to what extent the behavioral phenomena that have been taken to suggest model-based learning might arise from some simpler approximation. In this article, we explore a family of algorithms that capture a range of such approximations and, we argue, provide a promising set of candidates for the neural foundations supporting such learning.

Our proposals are motivated by multiple suggestive, but also somewhat counterintuitive, lines of evidence, which suggest that the dopamine system and its key targets are implicated not just in the model-free behaviors that theory endows them with, but also in the more flexible choice adjustments that seem to reflect model-based learning [5–11]. This is perplexing for the standard account because typical model-based algorithms such as value iteration do not make use of a TD prediction error for long-run reward of the sort associated with dopamine. Instead, they store internal variables (specifically, predictions about immediate “one-step” rather than long-run consequences of actions), which require different update rules and error signals [12, 13]. Additionally, at choice time, such algorithms require computations that are structurally different than those typically prescribed to the dopaminergic-striatal circuitry [14].

In this article, we revisit and extend the *successor representation* (SR) [15,16; see also 17-21], a predictive state representation that can endow TD learning with some aspects of model-based value computation, such as flexible adjustment following reward devaluation. That the SR, when combined with TD learning, can produce such flexible behavior makes it a promising candidate for a neural foundation for model-based learning, which (because it is built on top of a TD foundation) can also explain dopaminergic involvement in model-based learning. However, this approach, in its original form, results in behavioral inflexibilities that could serve as empirical signatures of it, but also make it inadequate for fully explaining organisms’ planning capacities. In particular, the SR simplifies planning by caching a set of intermediate quantities, expectations about cumulative future state occupancies. For this reason, unlike model-based learning, it is incapable of solving many tasks that require adjusting to changes in contingencies between actions and their long-run consequences (e.g. [22,23]).

In this article, we explore a family of algorithms based around the SR, and introduce two new variants that mitigate its limitations. In particular, we examine algorithms in which the SR is updated either by computation at choice time or off-line by replayed experience, both of which help to ameliorate its problems with caching. These approaches can each account for human and animal behavior in a wide range of planning tasks, suggest connections with other models of learning and dopamine, and make empirically testable predictions. Overall, these approaches represent a family of plausible and computationally efficient hypothetical mechanisms for the full range of flexible behaviors associated with model-based learning, and could provide a clear theoretical foundation for future experimental work.

The article is organized as follows. In the remainder of this introduction, we review the formalism of reinforcement learning in a Markov decision process (MDP) and use this framework to delineate the problem of model-based flexibility arising from model-free circuitry and elucidate how the SR offers a potential solution to this problem. In the results section, we use simulations to demonstrate the precise behavioral limitations induced by computing values using the SR, as originally described, and evaluate these limitations with respect to the behavioral literature. We then introduce two new versions of the SR that progressively mitigate these limitations, and again simulate their expected consequences in terms of flexible or inflexible choice behavior.

### Formalism: Model-based and model-free reinforcement learning

Here, we briefly review the formalism of reinforcement learning in a Markov decision process (MDP), which provides the foundation for our simulations (see [22] or [23] for a fuller presentation).

An MDP is defined by a set of states, a set of actions, a reward function *R*(*s*, *a*)over state/action pairs, and a state transition distribution, *P*(*s*′ |*s*, *a*), where *a* denotes the chosen action. States and rewards occur sequentially according to these one-step functions, driven by a series of actions; the goal is to learn to choose a probabilistic policy over actions, denoted by *π*, that maximizes the value function, *V*^*π*^ (*s*), defined as the expected cumulative discounted reward:

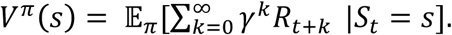

Here, *γ* is a parameter controlling temporal discounting. The value function can also be defined recursively as the sum of the immediate reward of the action chosen in that state, *R*(*s*, *a*), and the value of its successor state *s*’, averaged over possible actions, *a*, and transitions that would occur if the agent chose according to *π*:

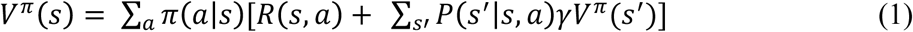

The value function under the optimal policy is given by:

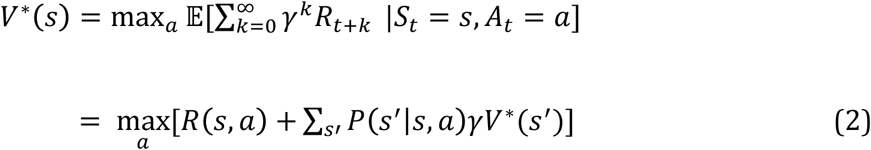

Knowledge of the value function can help to guide choices. For instance, we can define the state-action value function as the value of choosing action *a* and following *π* thereafter:

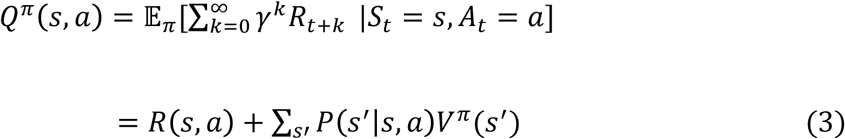

Then at any state one could choose the action that maximizes *Q*^*π*^ (*s*, *a*). (Formally this defines a new policy, which is as good or better than the baseline policy *π*; analogously, equation 2 can be used to define the optimal state-action value function, the maximization of which selects optimal actions.) Note that it is possible to write a recursive definition for *Q* in the same manner as equation 1, and work directly with the state-action values, rather than deriving them indirectly from *V*.

For expositional simplicity, in this article, we work instead with *V* wherever possible (mainly because this is easier to depict visually, and simplifies the notation), and accordingly we assume in our simulations that the agent derives *Q* using equation 3 for guiding choices. To be concrete, in a spatial “gridworld” task of the sort we simulate, this amounts to computing a value function *V* over locations *s*, and using it to derive *Q* (the value of actions *a* heading in each of the four cardinal directions) by examining *V* for each adjacent state. Although this simplifies bookkeeping for this class of tasks, *this is not intended as a substantive claim*. Indeed, the last algorithm we propose will work directly with *Q* values, and the others can easily be re-expressed in this form.

The problem of reinforcement learning is then reduced to learning to predict the value function *V*^*π*^ (*s*) or *V*\*(*s*). There are two main families of approaches. Model-based algorithms learn to estimate the one-step transition and reward functions, *P*(*s* ′ |*s*, *a*) and *R*(*s*, *a*), from which it is possible to compute *V*\* (or *Q*\*) using equation 2. This typically involves unrolling the recursion in equation 2 into a series of nested sums, an algorithm known as value iteration. The alternative, model-free, approach exemplified by TD learning bypasses estimating the one-step model. Instead, it directly updates a cached estimate of the value function itself. In particular, following a transition *s* → *s* ′ initiated by action *a*, a reward prediction error, *δ*, is calculated and used to update *V*(*s*):

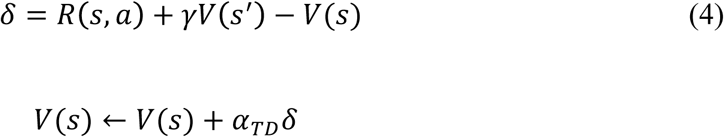

where *α*_*TD*_ is a learning rate parameter.

The TD update rule is derived from the recursion in equation 1: each step iteratively pushes the left hand side of the equation, *V*(*s*), closer to *R*(*s*, *a*)+ *γV*(*s*′), which is a one-sample estimate of the right hand side.

Finally, analogous sample-based updates may also be conducted offline (e.g., between steps of actual, “online” experience). This is a key insight of Sutton’s Dyna architecture [21] (see also [24]). The approach, like TD, caches estimates of *V*(*s*). Here TD learning is supplemented by additional offline updates. Specifically, samples consisting of a transition and reward triggered by a state-action (*s*, *a*, *r*, *s*’) are generated either from a learned one-step model’s probability distributions *P*(*s*′ |*s*, *a*) and *R*(*s*, *a*), or instead simply replayed, model-free, from stored episodes of previously experienced transitions. For each sample, *V*(*s*) is then updated according to equation (4). Given sufficient iterations of sampling between steps of real experience, this approach can substitute for explicit value iteration and produce estimates at each step comparable to model-based approaches that more directly solve equations 1 or 2.

A further distinction, which will become important later, is that between *on-policy* methods, based on equation 1, and *off-policy* methods, based on equation 2. On-policy methods estimate a policy-dependent value function *V*^*π*^(*s*), whereas off-policy methods directly estimate the optimal value function *V*\*(*s*). Typically, model-based methods are off-policy (since having learned a one-step model it is possible to use equation 2 to directly compute the optimal policy); whereas different TD learning variants can be either on- or off-policy.

### Model-free learning in the brain

Due to the similarity between the phasic responses of midbrain dopamine neurons, and the TD prediction error *δ* (equation 4), it has long been suggested that this system implements TD learning [2,3]. More specifically (e.g. [4]; Fig 1a) it has been suggested that values *V* or *Q* are associated with the firing of medium spiny neurons in striatum [28,29], as a function of an input state (or state-action) representation carried by their afferent neurons in frontal cortex, and that learning of the value function is driven by dopamine-mediated adjustment of the cortico-striatal synapses connecting these neurons. Selection among these striatal value representations would then drive action choice. Although not entirely uncontroversial, a great deal of evidence about dopamine and its targets supports this hypothesis (see [25,30] for fuller reviews).

**Fig. 1.**
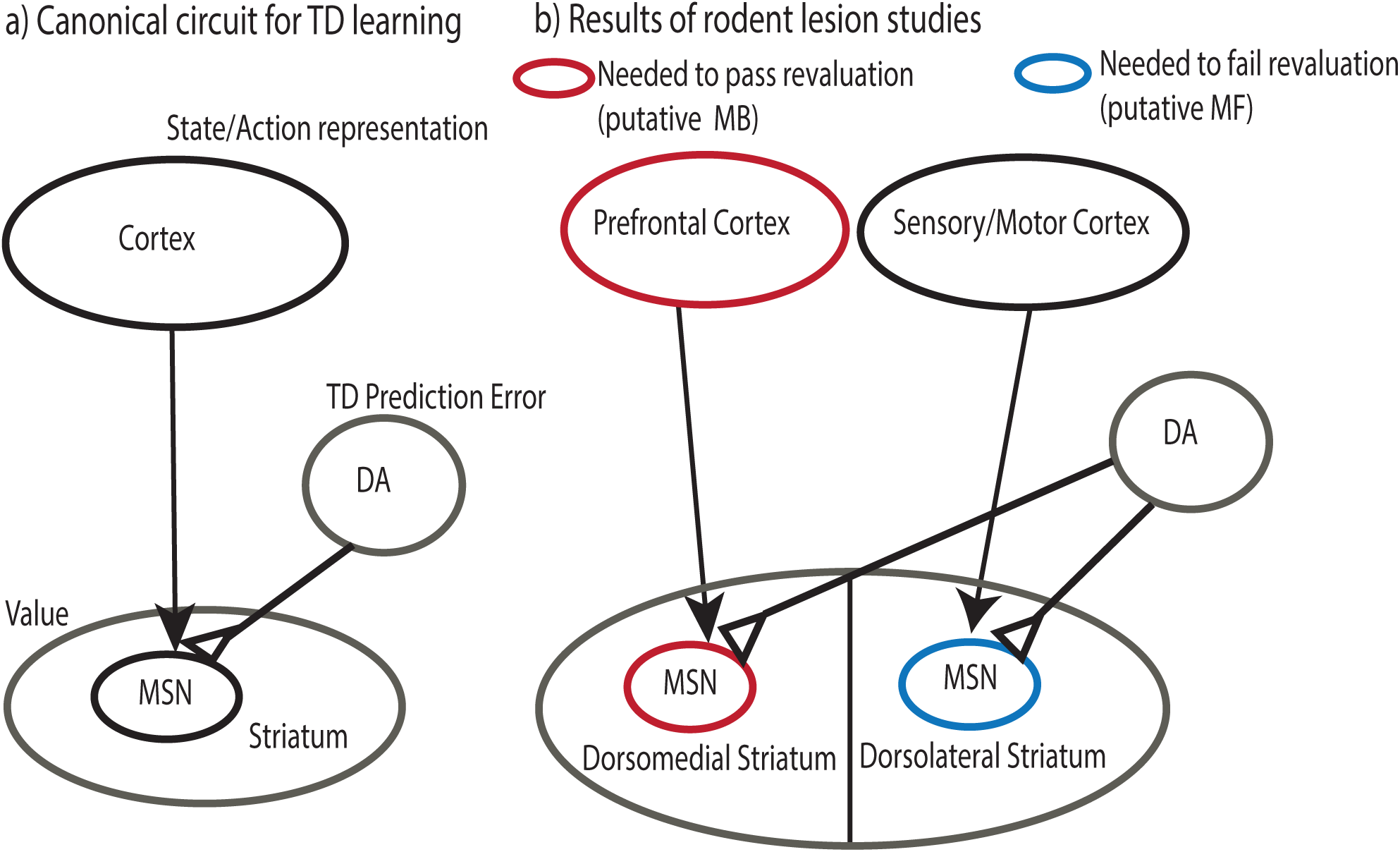
Cortico-striatal loops and reinforcement learning. a) Canonical circuit for TD learning. A dopaminergic prediction error, signaled in substantia nigra pars compacta and ventral tegmental area, updates the value of cortically represented states and actions by modifying cortico-striatal synapses. Depending on value, represented in striatal medium spiny neurons (MSN), actions are passed through to basal-ganglia action systems. b) Results of rodent lesion studies. Lesions to a cortico-striatal loop passing through dorsomedial (DM) striatum prevent flexibly adjusting behavior following reward devaluation. This area receives input from ventromedial prefrontal cortex and projects, via globus pallidus, to dorsomedial nucleus of the thalamus. This loop is generally thought to implement model-based learning [31]. Lesions to cortico-striatal loop passing through dorsolateral (DL) striatum cause animals to maintain ability to flexibly adjust behavior following devaluation, despite over-training. This area receives input from sensory and motor areas of cortex and projects, via globus pallidus, to posterior nucleus of the thalamus. This loop is generally thought to implement model-free learning [31]. In addition to receiving similar dopaminergic innervation from substantia nigra pars compacta (SnC), such loops are famously thought to be homologous to one another.

Such theories provide a neural implementation of Thorndike’s early law of effect, the reinforcement principle according to which rewarded actions (here, those paired with positive prediction error) tend to be repeated [32]. However, the hypothesis that animals or humans rely exclusively on this principle to make decisions has long been known to be false, as demonstrated by a line of learning experiments whose basic logic traces to rodent spatial navigation experiments by Tolman [22] (for modern variants, see [6,27,33–35]).

To facilitate simulation and analysis, we here frame the logic of these experiments in terms of “grid-world” spatial MDPs. When viewed as MDPs, Tolman’s experiments can be divided into two categories, which require subjects to adjust to either of two different sorts of local changes in the underlying MDP. Experience with these changes is staged so as to reveal whether they are relying on cached values or recomputing them from a representation of the full MDP.

Accordingly, revaluation tasks, such as latent learning, reward devaluation, and sensory preconditioning, examine whether animals appropriately adjust behavior following changes in *R*(*s*, *a*), such as a newly introduced reward (Fig 2a). Analogously, contingency change (e.g., detour or contingency degradation) tasks examine whether animals appropriately adjust behavior following changes in *P*(*s*′ |*s*, *a*), such as a blocked passageway (Fig 2b). Model-free RL is insensitive to these manipulations because it caches cumulative expected rewards and requires additional learning to update the stored *V*. Conversely, model-based RL, which uses the one-step model directly to compute *V* at decision time, reacts immediately and flexibly to any experience that affects it. Note that the difference in behavior on these types of tasks predicted by the algorithms is categorical, and not a question of degree or learning speed. In particular, because of the representations they learn and update, model-based algorithms can make the correct choice following these manipulations without any further retraining (i.e. so long as they learn locally about the new contingency or value, they can immediately make appropriate choices in distal parts of the state space), whereas model-free algorithms cannot (in general, they must first experience trajectories starting from the test state and leading to the state with the changed value or transition contingency). Animals sometimes fail to correctly update behavior following revaluations, consistent with inflexible, model-free caching schemes [36]. However, findings that in other circumstances animals can indeed flexibly adapt their behavior following such manipulations (without any further retraining – e.g. tested on the very first trial, or without feedback) has long been interpreted as evidence for their use of internal models, as in model-based RL or similar methods [1,22,37]. A key goal of this article is to interrogate this assumption, and to consider neural mechanisms that, despite falling short of full model-based RL, might support such behavioral flexibility.

**Fig. 2.**
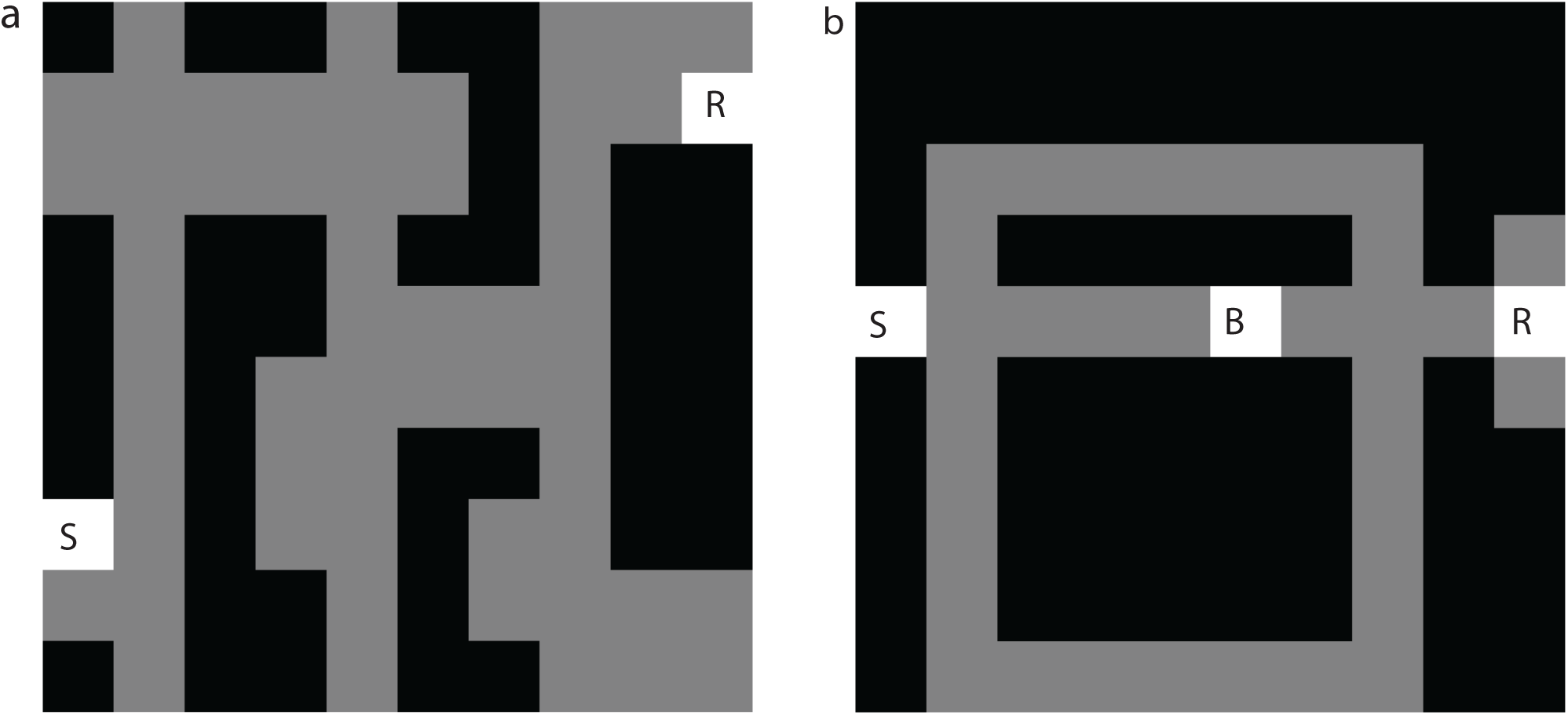
Grid-world representation of Tolman’s tasks. Dark grey positions represent maze boundaries. Light grey positions represent maze hallways. a) Latent learning: following a period of randomly exploring the maze (starting from S) the agent is notified that reward has been placed in position R. We examine whether the agent’s policy immediately updates to reflect the shortest path from S to R. b) Detour: after the agent learns to use the shortest path to reach a reward state R from state S, a barrier is placed in state B. After the agent is notified that state B is no longer accessible from its neighboring state, we examine whether its policy immediately updates to reflect the new shortest path to R from S.

### The puzzle of model-based learning and its neural substrates

A further set of rodent lesion studies have used reward devaluation tasks to suggest that apparently model-based and model-free behaviors (i.e., behavior that is either flexible or insensitive following reward devaluation) depend on dissociable sets of brain areas (e.g. [5,38]; Fig 1b). This led to the hypothesis (e.g., [1,31,39]) that these two forms of reinforcement learning depend on competing systems in the brain—the dopaminergic TD system previously described, plus a second – less clearly understood – circuit supporting model-based behavior. But how is this latter computation carried out in the brain?

A number of fairly abstract theories have been based around explicit computation of the state-action value based on some form of equation 3, e.g. by learning an estimate of the one-step transition function, *P*(*s*’|*s*, *a*) and the immediate reward function *R*(*s*, *a*) and using them iteratively to compute the future value by tree search, value iteration, or Bayesian inference [1,39–41]. These theories have not spoken in detail about the neural implementation of these computations, but an accompanying presumption has been that the model-based system does not rely on a dopaminergic prediction error signal. This is because the TD prediction error of equation 4 (for *γ* > 0, which is the parameter regime needed to explain phasic dopamine’s signature responses to the anticipation as well as receipt of reward [42]) is specifically useful for directly learning long-run cumulative values *V*. In contrast, the idea of model-based learning is to *derive* these values iteratively by stringing together short-term predictions from a learned one-step model [12,43]. Note that the prediction error normally thought to be reported by dopamine neurons is not appropriate here: the prediction error signal for updating the *immediate* reward model *R*(*s*, *a*) is like equation 4 but with *γ* = 0, which is not consistent with anticipatory phasic dopamine responses. (However, correlates of prediction errors for *γ* = 0 have been observed using human fMRI [44]). Furthermore, the hypothesized process of adding these rewards up over anticipated trajectories at choice time, such as by value iteration or tree search, has no counterpart in model-free choice. Instead, learning from anticipatory TD errors stores complete long-run values (e.g., in corticostriatal synapses), requiring no further computation at choice time.

However, neither the rodent lesion data nor another body of work studying the neural correlates of model-based learning in humans suggests such a clean differentiation between the dopaminergic-striatal circuit (supposed to support TD) and some other presumably non-dopaminergic substrate for model-based learning. Instead, lesions suggest each type of learning is supported by a different subregion of striatum, together with connected regions of cortex (Fig 1b) and basal ganglia. This suggests that putatively model-based and model-free systems may map onto adjacent but structurally parallel cortico-basal ganglionic loops [45], thus perhaps involving analogous (striatal) computations operating over distinct (cortical) input representations [46].

Also contrary to a strict division between systems, both dorsomedial and dorsolateral striatal territories have similar interrelationships with dopamine [47], though the involvement of their differential dopaminergic input in devaluation sensitivity has not been completely assessed [48]. Research on humans’ learning in a two-step MDP (which has similar logic to devaluation studies) supports the causal involvement of dopamine in model-based learning [7–10]. Furthermore, dopaminergic recordings in rodents [11] (though see [37]), and neuroimaging of prediction error signals in human striatum [6] suggest that these signals integrate model-based evaluations.

Altogether, the research reviewed here supports the idea that model-based evaluations are at least partly supported by the same sort of dopaminergic-striatal circuit thought to support TD learning, though perhaps operating in separate cortico-striatal loops. This suggestion, if true, provides strong hints about the neural basis of model-based behavior. However, for the reasons discussed above, this also seems puzzlingly inconsistent with the abstract, textbook [24] picture of model-based learning by equation 3.

Several more neurally explicit theories of some aspects of model-based computation have been advanced, which go some way toward resolving this tension by incorporating a dopaminergic component. Doya [49] introduced a circuit by which projections via the cerebellum perform one step of forward state prediction, which activates a dopaminergic prediction error for the anticipated state. The candidate action can then be accepted or rejected by thresholding this anticipated prediction error against some aspiration level. It is unclear, however, how this one-step, serial approach can be generalized to tasks involving stochastic state transitions, direct comparison between multiple competing actions, or rewards accumulated over multiple steps (as in tasks like [50]).

A similar idea has arisen from recordings in spatial tasks, where the firing of place cells along trajectories ahead of the animal suggests a hippocampal basis for a similar (though multi-step) state anticipation process, potentially driving evaluation of these candidate states using learned reward values in ventral striatum [51]. It is, however, unclear how this activity fits into a larger circuit for accumulation of these evaluations and comparison between options.

Finally, another candidate approach is based on the Dyna framework discussed above. In this case, model-generated experience can be played back “off-line,” e.g. between trials or during rest. These ersatz experiences can, in turn, drive dopaminergic prediction errors and updating of striatal *Q* values using the same mechanisms as real experience. As noted above, given sufficient off-line replay, this can achieve the same effect as model-based planning; in particular, it can update *Q* values following revaluation and other manipulations [27,52]. However, without a more traditional “on-line” planning component, this approach degrades (to that of basic, model-free *Q* learning) when there is insufficient time or resources for off-line replay, and when truly novel situations are encountered [27].

Here we propose and analyze a different family of approaches to these problems, which relate to the above proposals in that they incorporate elements of upstream predictive input to ventral striatum, and also of a different and more-flexible approach to offline updates. The proposed approach, based on the SR, builds even more directly on the standard TD learning model of dopaminergic-striatal circuitry.

### The successor representation

The research reviewed above suggests that flexible, seemingly model-based choices may be accomplished using computations that are homologous to those used in model-free RL. How can this be? In fact, it is known that evaluations with some features of model-based learning can result from TD learning over a different input representation. As shown by Dayan [15], equation 1 can reformulated as:

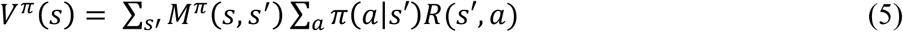

Here, *M*^*π*^ is a matrix of expected cumulative discounted *future state occupancies*, measuring the cumulative time expected to be spent in each future state *s*′, if one were to start in some state *s* and follow policy *π* (Fig 3):

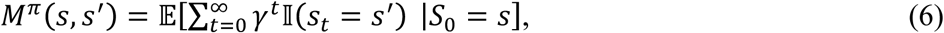

where 𝕀(·) = 1 if its argument is true and 0 otherwise. Thus, this form rearranges the expectation over future trajectories in equation 1 by first computing expected occupancies for each state, then summing rewards obtained, via actions, in each state over these.

*M*^*π*^ can also be used as a set of basis functions for TD learning of values. Specifically, we represent each state using a vector of features given by the corresponding row of *M* (Fig 3), i.e. by the future occupancies expected for each state *s*′. Then we approximate *V* ^*π*^ *s* by some weighted combination of these features:

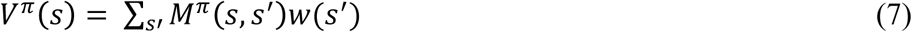

**Fig. 3.**
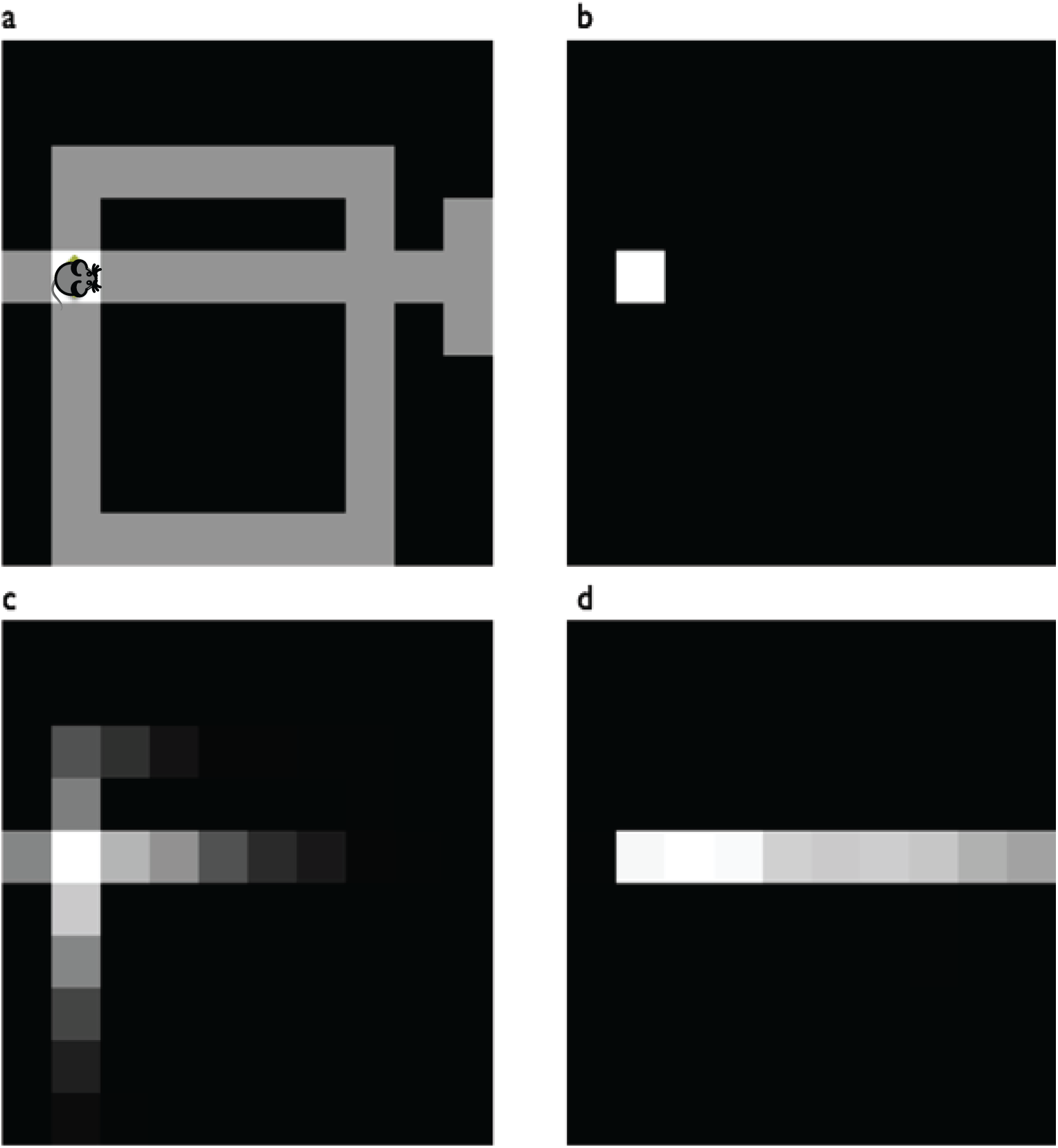
Example state representations. a) Agent position (rodent image) in a maze whose hallways are indicated by grey. b) Punctate representation of the agent’s current state. Model-free behavior results from TD computation applied to this representation c,d) Possible successor representations of agent’s state. Model-based behavior may result from TD applied to this type of representation. The successor representation depends on the action selection policy the agent is expected to follow in future states. The figures show the representation of the current state under a random policy (c) versus a policy favoring rightward moves (d).

Comparing equations 5 and 7 demonstrates this approximation will be correct when the weight *w*(*s*′) for each successor state corresponds to its one-step reward, averaged over actions in *s*′, ∑_*a*_ *π(a*|*s*′)*R*(*s*′, *a*). One way to learn these weights is using standard TD learning (adapted for linear function approximation rather than the special case of a punctate state representation). In particular, following a transition *s* → *s*′, each index *i* of *w* is updated:

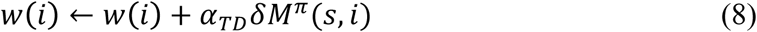

Here, *δ* is defined as in equation 4. Note that in the algorithms discussed below, the agent must estimate the successor matrix *M*^*π*^ from experience. If the feature matrix *M*^*π*^ were known and static, a simpler alternative to equation (8) for *w* would be to learn the one-step rewards by a delta rule on the immediate reward *R*. Since the successor representation is just a particular case of a linear feature vector for TD learning, the advantage of learning weights by the TD rule of equation 8 is that weights learned this way will estimate value *V*^*π*^ for any feature matrix *M*, such as estimates of the successor matrix *M*^*π*^ prior to convergence (Supplementary Figure 1).

Altogether, this algorithm suggests a strategy for providing different inputs into a common dopaminergic/TD learning stage to produce different sorts of value predictions (see also [16]). In particular, whereas model-free valuation may arise from TD mapping of a punctate representation of the current state (Fig 3b) in sensory and motor cortex to values in dorsolateral striatum (Fig 1b), at least some aspects of model-based valuation may arise by analogous TD mapping of the successor representation (Fig 3c,d) in prefrontal cortex or hippocampus to values in dorsomedial striatum (Fig 1b). This is possible because the successor matrix *M* has a predictive aspect reflecting knowledge of the state transitions *P*(*s*′ |*s*, *a*), at least in terms of aggregate occupancy, separate from the state/action rewards *R*(*s*, *a*).

This approach may thus offer a solution to how flexible, seemingly model-based choices can be implemented, and indeed can arise from the same dopaminergic-striatal circuitry that carries out model-free TD learning.

What remains to be shown is whether algorithms based on this strategy – applying the SR as input to TD learning – can produce the full range of model-based behaviors. In the remainder of this paper, we simulate the behavior of such algorithms to explore this question.

To simulate learning using the SR, we need to also simulate how the successor matrix *M*^*π*^ is itself produced from experience. *M*^*π*^ can be defined through a recursive equation that is directly analogous to equations 1 and 2:

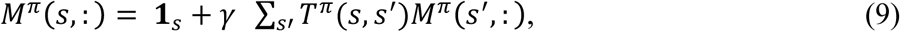

where **1**_*s*_ is the vector of all zeros except for a 1 in the sth position and *T*^*π*^ is the one-step state transition matrix that is dependent on *π*, *T*^*π*^(*s*, *s*′)=∑_*a*_*π* (*a* |*s*) *P*(*s*′|*s*, *a*).

Similar to how approaches to estimating *V* are derived from equations 1 and 2, one could derive analogous approaches to estimating *M*^*π*^ from equation 9. Specifically, one could utilize a “model-based” approach that would learn *T*^*π*^ and use it iteratively to derive a solution for *M*^*π*^. Alternatively, a TD learning approach could be taken to learn *M*^*π*^ directly, without use of a one-step model *T*^*π*^. (This approach is analogous to model-free TD methods for learning *V*, though it is arguably not really model-free since *M*^*π*^ is itself a sort of long-run transition model.) This TD learning approach would cache rows of M and update them after transitioning from their corresponding states, by moving the cached row closer to a one-sample estimate of the right hand side of equation 9. Lastly, such TD updates could also occur offline, using simulated or previously experienced samples. This approach for learning *M*^*π*^ would be comparable to the Dyna approach for learning *V*. The three models we consider below correspond to these three different possibilities.

Finally, note that SR-based algorithms have favorable computational properties; in particular, at choice time, given *M*^*π*^ (e.g. if it is learned and cached rather than computed from a one-step model), SR can compute values *V*^*π*^ with a single dot product (e.g., a single layer of a linear neural network, equation 7), analogous to model-free TD algorithms. This is in contrast to the multiple steps of iterative computation required at choice time for computing value via equation 1 in standard model-based approaches. This comes at the cost of storing the successor matrix *M*^*π*^: if S is the number of states in the task, the SR matrix has a number of entries equal to *S*^2^. Such entries of *M*^*π*^can be stored as the (all-to-all) set of weights from a single layer of a neural network mapping input states to their successor representation.

## Results

In the following sections, we explore the behavioral consequences of each of these strategies. We structure the results as follows. For each learning method, we first present the algorithm. Then we present the results of simulations using that algorithm. The purpose of simulations is to verify our qualitative reasoning about the behavior of the algorithm and illustrate how the algorithm’s behavior compares to that of model-based dynamic programming methods. These simulations also suggest experiments that could be used to identify whether an animal or human were planning using such a strategy. Each task that we simulate is designed to be a categorical test the algorithm. Following some change in the task to which the agent must respond, some of the algorithms can arrive at the correct decision without additional experience, but other algorithms cannot. Such failures are due to the computational properties of the algorithms themselves and are thus parameter-independent. To ensure that this is the case, for each simulation presented in the results, we have verified that the qualitative result can be observed robustly under a wide range of parameter settings. In general, there are parameter settings under which models, which are demonstrated below to succeed in a given task, can be made to fail it. However, there are *no parameter settings* under which a model that is shown below to fail a given task will pass it (Supplementary Table 1).

For each algorithm, we discuss its biological plausibility as well as how that algorithm’s performance lines up with that of animals.

### Algorithm 1: The Original Successor Representation (SR-TD)

The original SR [15] (which we call SR-TD) constructed the future state occupancy predictions *M*^*π*^ using a TD learning approach. This approach caches rows of *M*^*π*^ and incrementally updates them after transitioning from their corresponding states. Specifically, following each state transition *s* → *s*′ each element of row *s* is updated as follows:

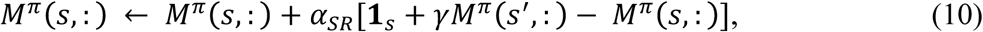

where **1**_*s*_ is the vector of all zeros except for a 1 in the sth position. *M*^*π*^(*s*,:) is used as input to another TD learning stage, this time to learn the weights *w* for predicting expected future value from the state occupancy vector. To simulate SR-TD, we have the agent learn *M*^*π*^and *w* in parallel, updating each (according to equations 10 and 8, respectively) at each transition; and sample actions according to an *ϵ*-greedy policy (see Methods).

### Simulation Results

*SR-TD can solve some reward revaluation tasks*. SR-TD is able to produce behavior analogous to model-based learning in some reward revaluation tasks that categorically defeat simple TD learning. To demonstrate this, we simulated the behavior of SR-TD in a grid-world version of Tolman’s *latent learning* task (Fig 2a,b). The agent first explores the grid-world randomly, during which it learns the successor matrix ***M*** ^*π*^ corresponding to a random policy. Next, it learns that reward is available at position R (importantly, by being placed repeatedly at R and receiving reward, but not experiencing trajectories leading there from any other location). This experience induces prediction errors that cause the agent to update its weight vector, ***w*** in the position corresponding to the rewarded state. Finally, in a test probe, we allow the agent to form values by multiplying its current version of ***M***^***π***^ with ***w***, and measure whether (immediately on the first test trial following reward training) those values would produce a policy reflective of the shortest path from position S to R.

Fig 4b shows SR-TD performance on a latent learning task: SR-TD can, without further learning, produce a new policy reflecting the shortest path to the rewarded location (Fig 4b). As a comparison to SR-TD’s performance, we also simulated the behavior of a simpler foil algorithm that represents model-free (or, actually, limited model-based) performance. This algorithm applied TD learning updates to non-predictive, punctate, state representations to estimate *V*. As with the SR, we permitted this algorithm to convert state values to state-action values by using a single-step of model-based look-ahead. Although this algorithm’s performance is representative of the failure of fully model-free algorithms at solving these revaluation tasks, we designed it to go beyond a vanilla model-free TD algorithm by allowing a single-step of model-based lookahead. This is analogous to a limited sort of model-based learning that has been suggested previously to be implemented by the basal ganglia and cerebellum [49]. Fig 4a shows that this algorithm cannot solve latent learning problems: it learns nothing about paths around the maze from the reward training, and would have to discover the path to the new reward from scratch by additional exploration.

**Fig. 4.**
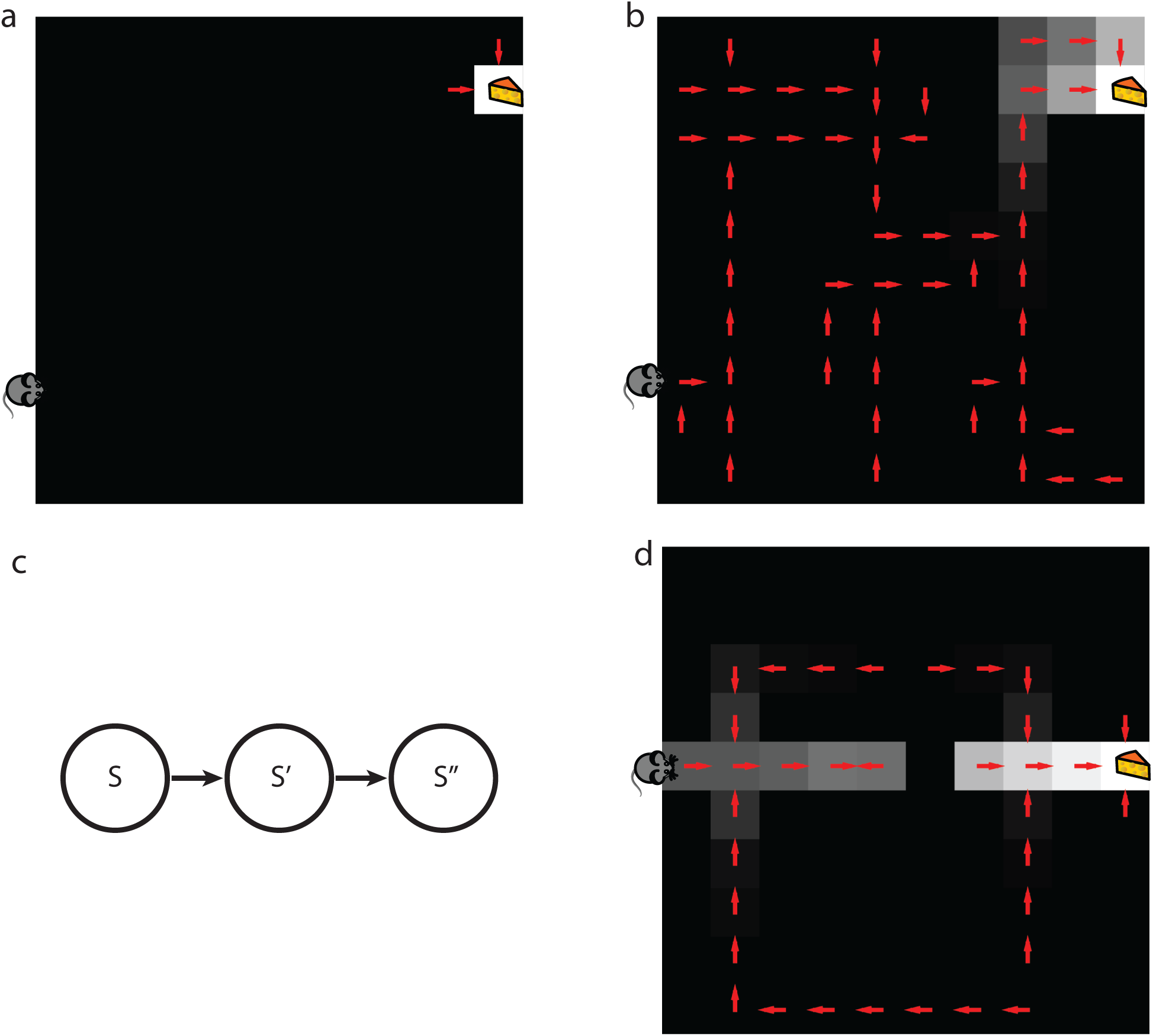
Behavior of SR-TD: a) One-step of model-based lookahead combined with TD learning applied to punctate representations cannot solve latent learning tasks. b) SR-TD can solve some latent learning tasks. For both a) and b) median value function (grayscale) and implied policy (arrows) are shown immediately after the agent learns about reward in latent learning task. c) SR-TD can only update predicted future state occupancies following direct experience with states and their multi-step successors. For instance, if SR-TD were to learn that s’’ no longer follows s’, it would not be able to infer that state s’’ no longer follows state s. Whether animals make this sort of inference is tested in the detour task. d) SR-TD cannot solve detour problems. Median value function (grayscale) and implied policy (arrows) are shown after SR-TD encounters barrier in detour task. SR-TD fails to update decision policy to reflect the new shortest path.

*SR-TD cannot solve transition revaluation tasks*. However, SR-TD is limited in its ability to react correctly to other seemingly similar manipulations. Because ***M***^***π***^ reflects long-run cumulative state occupancies, rather than the individual one-step transition distribution, ***P***(***s***’ **|*s***, ***a***), SR-TD cannot adjust its valuations to local changes in the transitions without first updating ***M*^*π*^**at different locations. This inflexibility prevents SR-TD from flexibly adjusting value estimates after learning about changes in transition structure (“transition revaluation”; Fig 4b). Consider a grid-world version of Tolman’s detour task (Figs 2b and 4d). Here, following an exploration period during which the agent is able to form an estimate of ***M*^*π*^** under a random policy, the agent is trained to seek reward at R, starting from S. Later, a blockade is introduced at B. Again, the agent is allowed to experience this change only *locally*, by repeatedly being dropped in the state next to the barrier, attempting the action that leads to it and learning that the states to the right are no longer accessible from this state. This experience causes the agent to update ***M*^*π*^** for the state immediately next to the barrier. However, despite this update, the rows of ***M*^*π*^** corresponding to the states that lie along a path between the start state and the state next to the barrier remain unchanged. From equation 10, it can be seen that these updates can only occur from direct experience, i.e., a series of new trajectories starting at these states that encounter the barricade. SR-TD fails to reduce the value of these states (Fig 4d), and thus would approach the barricade rather than taking a detour on the first visit back to S. As shown in supplemental materials, the depth-limited model-free algorithm also fails this test (Supplementary Table 1). A fully model-based algorithm (not shown) does make the correct choice in this case.

### Interim Discussion

#### Biological Plausibility

The reward learning stage of this rule (learning weights ***w*** to map ***M*^*π*^ (*s***,:) to ***V***^***π***^ (***s***)) is the standard dopaminergic TD rule, equation 8, operating over a new input. The update rule for that input, ***M*^*π*^(*s***,: ), is also based on a TD learning rule, but here applied to learning to predict cumulative future state occupancies. This uses a vector-valued error signal to update an entire row of ***M*^*π*^** at each step. Crucially, despite the functional similarity between this rule and the TD update prescribed to dopamine, we do not suggest that dopamine carries this second error signal. Neurally, this sort of learning might, instead, be implemented using Hebbian associative learning between adjacent consecutive states [53], with decaying eligibility traces (like TD(1)) to capture longer-run dependencies. Lastly, although we have defined the successor representation over tabular representations of states, is also possible to combine the SR with function approximation and distributed representations in order to reduce its dimensionality [21,54].

#### Behavioral Adequacy

SR-TD is capable of solving some reward revaluation experiments. For similar reasons, SR-TD can solve sensory preconditioning (e.g. [35]) and reward devaluation tasks (e.g. [6,27,33,34]), both of which turn on an analogous ability to update behavior when state transition probabilities are held constant but reward values are changed. Evidence for model-based behavior in animals and humans has typically come from these types of tasks, suggesting that SR-TD could underlie a good proportion of behavior considered to be model-based. However, SR-TD is incapable of solving seemingly analogous tasks that require replanning under a transition rather than a reward change. Because there is at least some evidence from the early literature [55] that animals can adapt correctly to such detour situations, we suggest that this inflexibility prevents SR-TD, on its own, from being a plausible mechanism for the full repertoire of model-based behavior.

##### Algorithm 2: Dynamic recomputation of the successor representation (SR-MB)

Here, we explore a novel “model-based” approach, SR-MB, for constructing the expected state occupancy vector *M*^*π*^ (*s*,:). SR-MB learns a one-step transition model, *T*^*π*^ and uses it, at decision time, to derive a solution to equation 9. One key constraint on a model-based implementation suggested by the data is that the computation should be staged in a way consistent with the architecture suggested by Fig 2a. Specifically, the TD architecture in Fig 2a suggests that, because the states are represented in cortex (or hippocampus) and weights (which capture information about rewards) and value are represented in downstream cortico-striatal synapses and medium spiny striatal neurons, information about *R*(*s*, *a*) and *V*(*s*) should not be used in the online construction of states. For the SR approach, this implies that *M* be constructed without using direct knowledge of *R*(*s*, *a*) or *V*(*s*). As we see below, this serial architecture – a cortical state-prediction stage providing input for a subcortical reward-prediction stage – if true, would impose interesting limitations on the resulting behavior.

To construct *M*^*π*^ (*s*,:), SR-MB first learns the one-step state transition matrix *T*^*π*^, implemented in our simulations through separate learning of *P* (*s*′ | *s*, *a*) as well as *π* (*a*|*s*), the agent’s previously expressed decision policy (see Methods). Prior to each decision, *T*^*π*^ is used to compute a solution to equation 9. This solution can be expressed in either of two forms. A given row, *s*, of *M* can be computed individually as the sum of n-step transition probabilities starting from state *s*:

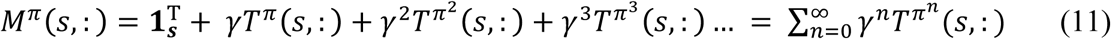

Alternatively, matrix inversion can be used to solve for the entire successor matrix at once:

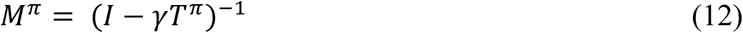

To implement SR-MB, we use equation 12. However, this is not a mechanistic commitment of the model, since equation 11 is equivalent.

Given *M*^*π*^, SR-MB learns the reward prediction weights *w* and forms *V* and *Q* values in the same way as SR-TD.

Note finally that this scheme is similar to solving equation 1 for on-policy values *V*^*π*^ by value iteration, except that the sums are rearranged to put state prediction upstream of reward prediction, as per equation 7 and in line with the neural architecture of Figure 2a. The max operator in equation 2 prevents a similar rearrangement that would allow this scheme to be used for off-policy optimal values *V*\* (equation 2), as discussed below. The restriction to on-policy values *V*^*π*^ is the major empirical signature of this version of the model.

### Simulation Results

#### SR-MB can solve transition revaluation tasks

Using an updated *T*^*π*^ to recompute *M*^*π*^ at decision time ensures that behavior is sensitive to changes in the transition structure. We demonstrate this by showing that unlike SR-TD, SR-MB successfully solves Tolman’s detour task in addition to latent learning. In the detour task, after being dropped in the state next to the barrier, SR-MB updates its estimate of *P*(*s*’|*sa*) for the *sa* leading into the barrier. This new *P*(*s*’|*sa*) is then combined with it’s estimate of *π* (*s*, *a*) (learned through prior experience including initial random exploration of the maze and then trials starting in S and ending in R) to compute *T*^*π*^. The row of *T*^*π*^ corresponding to the state next to the barrier at this point now no longer predicts transitioning into the barrier state. When the agent then recomputes *M*^*π*^ by equation 12, using the updated *T*^*π*^, rows of *M*^*π*^ corresponding to the path between the start state and the barrier no longer predict future occupancy of states on the other side of the barrier. When *M*^*π*^is then used to compute *V*^*π*^, the values immediately (on the first test trial after the barrier is encountered) result in a policy reflective of the new shortest path (Fig 5a.)

**Fig. 5.**
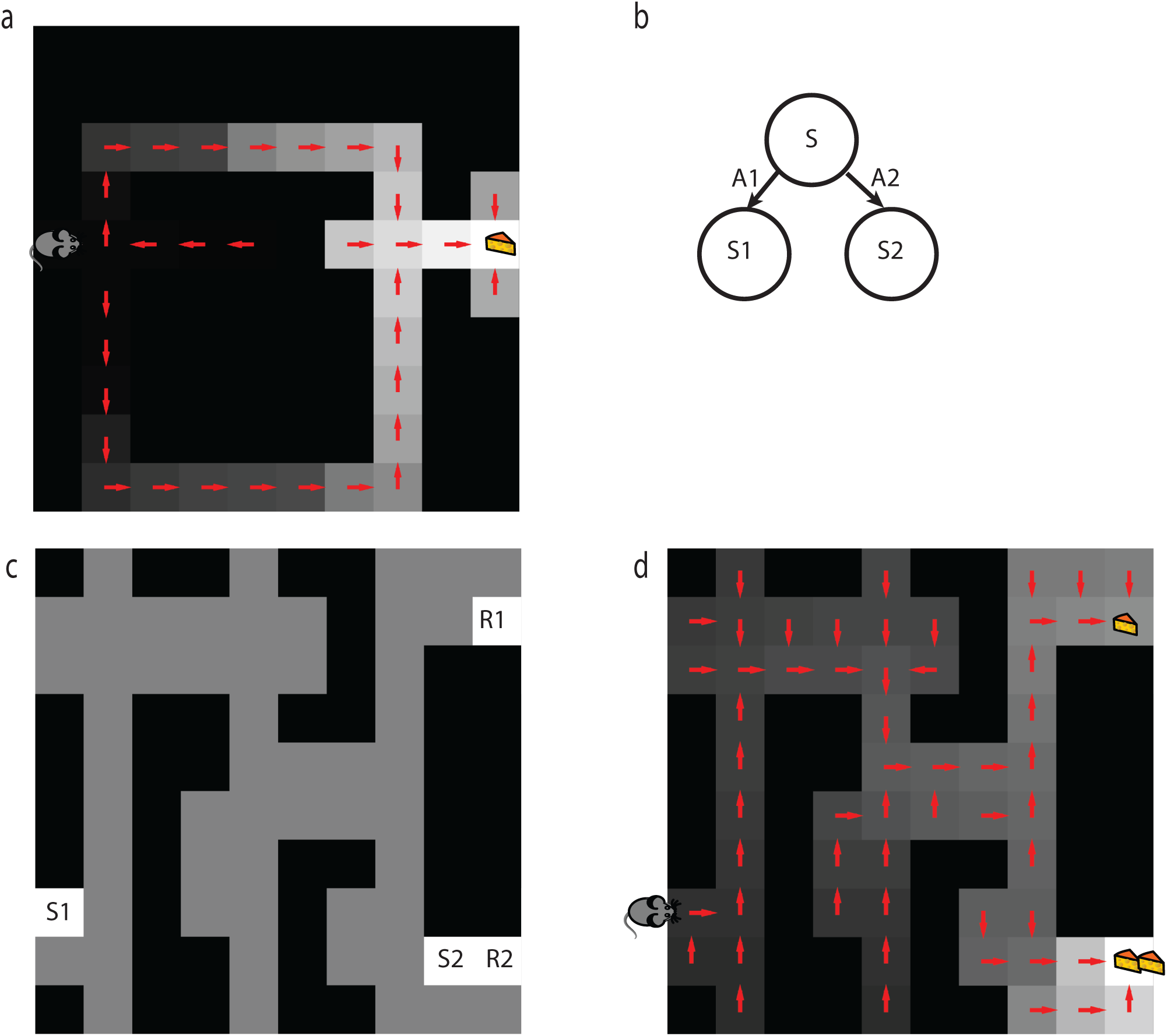
Behavior of SR-MB. a) SR-MB can solve the detour task. Median value function (grayscale) and implied policy (arrows) after SR-MB encounters barrier. b) SR-MB determines successor states relative to a cached policy. If SR-MB learned from previous behavior that it will always select action a1, the value of s would become insensitive to changes in reward at s2’. C) Novel “policy” revaluation task. After a phase of random exploration, we place a reward in location R1. The agent completes a series of trials that alternatively start from locations S1 and S2 and end when R1 is reached. We then place a larger reward in location R2 and record the agent’s value function and implied policy upon encountering it. d) SR-MB cannot solve the novel “policy” revaluation task. Median value function and implied policy recorded immediately after SR-MB learns about reward placed in location R2. Notice that if the agent were to start from location S1, its policy would suboptimally lead it to the smaller reward at R1.

#### SR-MB is limited by policy dependence

SR-MB is able to solve both tasks that have been used as examples of planning in animals and humans. We thus sought to determine whether there were any tasks, perhaps not yet explored in the empirical literature, that could differentiate it from approaches that utilize “full” model-based value iteration. A key feature of SR-MB, as well as SR-TD, is that it computes ***M*^*π*^** with respect to a policy ***π***. For SR-MB, ***T*^*π*^** is computed using ***π***(***a*** |***s***), which is learned through observation of previous actions. Because ***M*^*π*^** is policy dependent, so are the value estimates that it produces. SR-TD and SR-MB are thus “on-policy” methods – their estimates of ***V***^***π***^ can be compared to the estimates of a traditional model-based approach used to solve equation 1. A key limitation of “on-policy” methods is that their value estimates are made with respect to the policy under which learning occurred. This is an important limitation, as we will see below, because new learning about parts of the MDP can moot the previously learned policy ***π*** (and hence invalidate the associated successor matrix and values).

For the successor representation strategy, this limitation could be bypassed if we could compute ***M***\* – an “off-policy” successor matrix based on the successor states expected under the *optimal* policy – which would, in turn provide the state input for solving for ***V***\*, the optimal long-run values. However, given the architectural constraints we have suggested, computing ***M***\* is not straightforward. In particular, defining equation 9 with respect to the optimal policy would require replacing ***T*^*π*^** with *T\**: the one-step transition matrix corresponding to the optimal policy, ***T***\*(***s***, ***s***’) = ***P***(***s***’|***s***, ***a***\*), where ***a***\* is the action in state ***s*** that maximizes future rewards. Computing ***a***\* online, however, would require access to ***R***(***s***, ***a***), which would violate the suggested serial staging of the computation, that ***M*** be constructed without using reward information. Policy dependence of value predictions thus defines both the major limitation as well as the empirical signature of SR-MB.

#### SR-MB cannot solve novel policy revaluation tasks

What are the behavioral implications of SR-MB estimating values using the policy that was expressed during learning? Consider a situation where state ***s***’ can be reached from state ***s*** using action ***a***, but SR-MB learned from past behavior that ***π***(***a***|***s***) is near 0 (Fig 5b). Then it will not include rewards at ***s***’ in the value of ***s***, even if separately learning (say, by visiting ***s***’ but not in a trajectory starting from ***s***) that ***s***’ is the most rewarding state reachable from ***s***. In other words, caching of the policy at ***s*** blinds the agent (without further exploration and relearning) to changes in the reward function that should change that policy. Value iteration based on equation 2 does not have this limitation because the max operation would look ahead to the reward at ***s***’ to determine whether it should be included.

These considerations suggest a novel revaluation task (Fig 5c). Here, SR-MB first performs many trials where R1 is rewarded, starting from both S1 as well as a new start position, S2. This experience causes the agent to learn a policy ***π***(***s***, ***a***) that reflects moves away from the bottom right corner of the maze. Next, a larger reward is introduced at R2. Despite having learned about this reward (by starting at R2 and updating ***w*** at the state corresponding to this location), because computation of ***M*^*π*^** utilizes ***T*^*π*^**, which reflects moves away from the bottom right corner of the maze, ***M*^*π*^** does not predict the future occupancy of state R2 from any state along a path starting at S1. Because of this, the values of these states do not update to include the newly updated parts of the weight vector corresponding to position R2. Thus, despite the higher reward in R2, the agent would choose to head toward R1 from S1, due to caching of the incorrect policy in ***T*^*π*^** (Fig 5d). Thus, this task defeats SR-MB (as well as, shown in supplemental materials, the depth limited model-free planners and SR-TD), though it can be solved by standard model-based learning using equation 2.

### Interim Discussion

#### Biological Plausibility

SR-MB requires the brain to compute *M*^*π*^(*s*,:) from *T* ^*π*^ (*s*,:) for a particular state s under evaluation. Although in our simulations, we used equation 12 to compute the entire successor matrix at once, this is not a mechanistic commitment of the model. For instance, recurrent neural networks offer a simple way to compute *M*^*π*^(*s*,:) based on spreading activation implementing equation 11. Consider a network with one node for each state and the weight between node *s* and node *s*’ set to *γT* ^*π*^ (*s*, *s*′). If node *s* is activated, then at each successive time step, the network activity will represent each successive component of the sum. Indeed, this model arose in the early connectionist literature [17].

Alternatively, it is also well established that recurrent neural networks can perform matrix inversion by relaxing to an attractor [56], making a computation based on equation 12 plausible as well.

A final mechanism for computing *M* from *T* would involve sampling transitions from *T* offline and using them to iteratively update *M* according to equation 10, the SR-TD update. In the following section, we explore how an approach based on this idea, when carried out over state-actions, can be used to solve the “off-policy” planning problem as well.

#### Behavioral Adequacy

SR-MB produces the two behaviors that are considered signatures of planning in the empirical literature: immediate adjustment of decision policy following learned changes in either reward structure (latent learning) or transition structure (detour problem). The novel policy revaluation task demonstrates that SR-MB still produces errors that could in principle be behaviorally detectable, but have not been exercised by standard experimental tasks [57].

##### Algorithm 3: Off-policy experience resampling (SR-Dyna)

Here we introduce a third approach towards solving equation 9, SR-Dyna, which can be compared to Sutton’s Dyna approach [26] for solving equations 1 and 2. Akin to how Dyna replays experienced transitions offline to update estimates of *V*(*s*), SR-Dyna replays experienced transitions to update the successor matrix. When this approach is combined with an ‘off-policy’ update rule, similar to Q learning, to update the successor matrix offline, it is capable of solving the off-policy planning problem. Utilizing this type of update, however, requires us to work with a state-action version of the successor representation, *H*, which can be used directly to form *Q* values [58,59]. The key idea here is to define future occupancy not over states but over state/action pairs, *sa*. Analogous to equation 3, *Q* ^*π*^ can then be expressed:

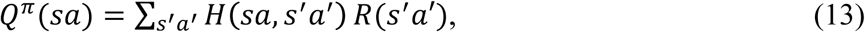

*H* is a matrix of expected cumulative discounted future state-action visitations, i.e. given that you are starting with state *s* and action *a*, the cumulative (discounted) expected number of times you will encounter each other state/action pair:

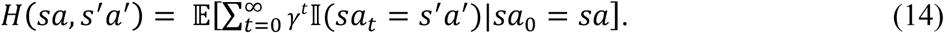

*H* can then be used as a linear basis for learning *Q*(*s*, *a*), using the SARSA TD algorithm to learn a weight for each column of *H*. In particular, when state-action *s*’*a*’ is performed after state action *sa*, a prediction error is calculated and used to update w:

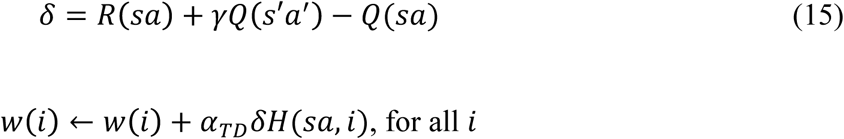

Like *M*, *H* can be defined recursively:

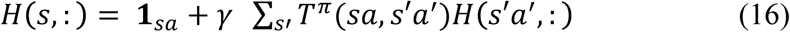

where *T* ^*π*^ is the one-step state-action transition matrix, *T* ^*π*^ (*sa*, *s*′*a* ′) = ∑_*s*′_ ∑_*a* ′_ *P*(*s*′ |s, *a*) *π*(*a* ′|*s*′). As with SR-TD, this recursion can be used to derive a TD-like update rule by which an estimate of *H* can be iteratively updated:

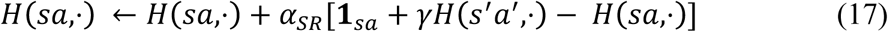

As with SR-MB, it is also possible to derive ***H*** from ***T*^*π*^ (*sa***, ***s***′ ***a***′) using an explicit “model-based” solution analogous to equation 9. However, here, we investigate the approach of updating ***H*** off-line (e.g., between trials or during rest periods) using replay of experienced trajectories (e.g. [60]). The key assumption we make is that this off-line replay can sequentially activate both the state and reward (cortical and basal ganglia) stages of Figure 1b, giving rise to an off-policy update of *H* with respect to the policy ***π***\* that is optimal given the current rewards. By comparison, as articulated above, we assumed such policy maximization was not possible when computing the successor representation ***M*** on-line for SR-MB, since this entire computation was supposed to happen in cortex at decision time, upstream of the striatal reward learning stage.

Following each transition, SR-Dyna stores the sample (*s*, *a*, *s*’). Then in between decisions, SR-Dyna randomly selects (with a recency weighted bias) *k* samples (with replacement). For each sample, it updates H as follows:

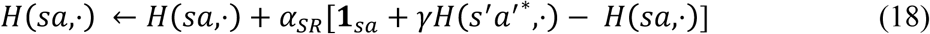

where

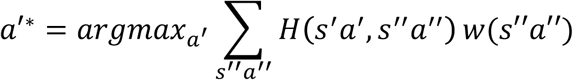

That is, when *H* updates from previously experienced samples, it performs an off-policy update using the best action it could have chosen, rather than the one it actually chose.

## Results

### SR-Dyna can solve policy revaluation tasks

Given sufficient sampling (large enough ***k***), this off-policy updating not only permits SR-Dyna to solve the detour task, but also to solve the novel policy revaluation task (Fig 6a). In the policy revaluation task, after the agent is introduced to the new reward in R2, and updates ***w***, we permit it to draw 10,000 random samples from memory and perform an update for each. With each sample drawn, SR-Dyna partially adjusts the predictions of future state occupancies from a given state action ***sa***, so that they become closer to the predictions of future state occupancies from ***s***’***a***\*, where ***a***\* is chosen with consideration of the newly updated ***w***. Such updates thus allow SR-Dyna to re-learn a new version of ***H*^*π*^**, corresponding to the policy that would result from repeated choices under the updated ***w***. Once updated, ***H*^*π*^** comes to reflect prediction of future states reflective of a policy that moves towards the new highest reward in the bottom right of the maze. When the updated ***H*^*π*^** is then used to compute new values (on the first test trial following the policy retraining), those values result in a policy that would bring the agent from S to the new highest reward in R2 along the shortest path. This simulation demonstrates that SR-Dyna can thus produce behavior identical to “full” model-based value iteration in this task (as well as the other revaluation tasks previously simulated, as shown below in Figure 6). However, it has the potential advantage that updating can take place fully off-line and thus offload computation to situations that may be ecologically convenient such as sleep or wakeful rest.

**Fig. 6.**
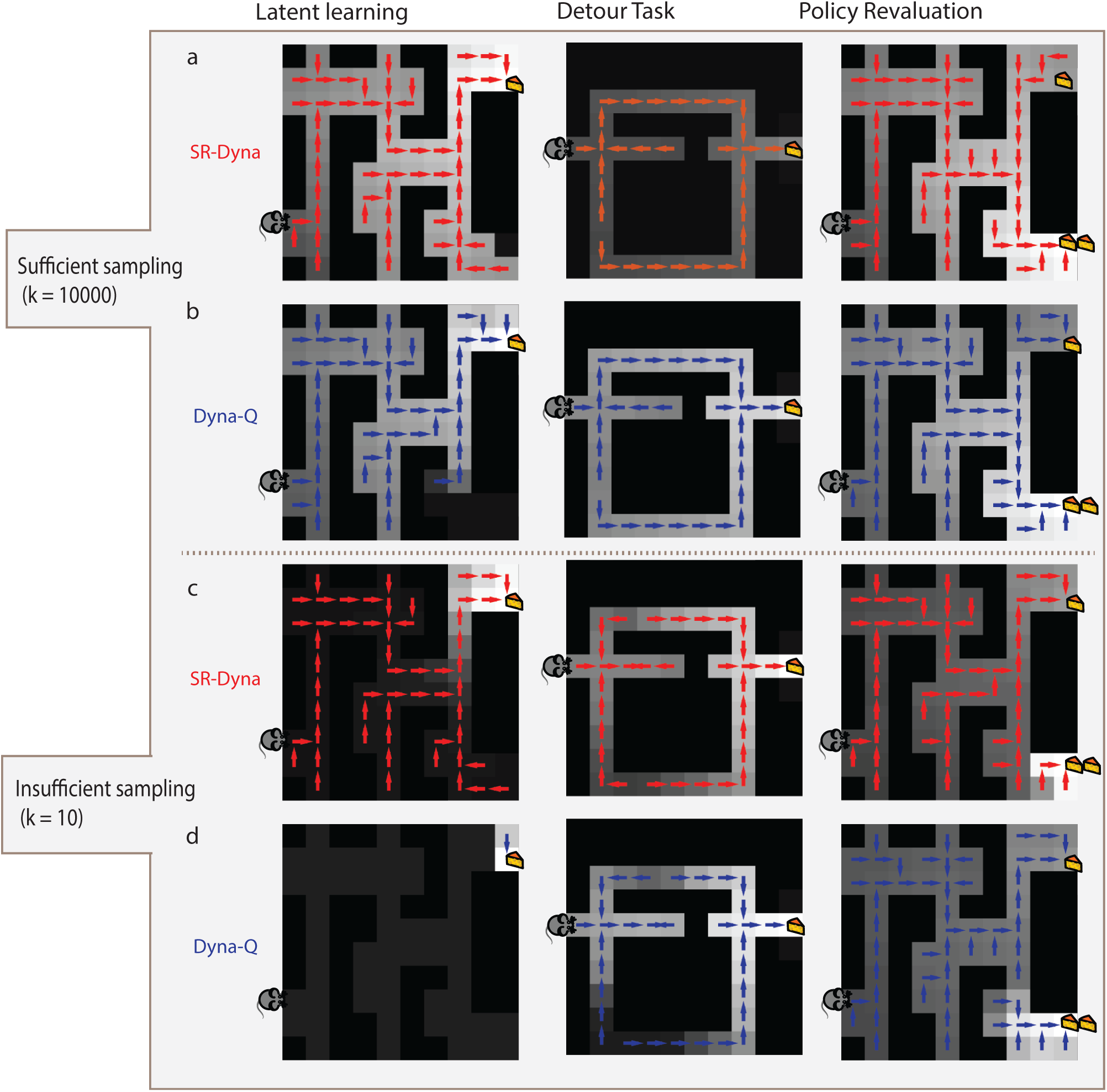
Comparison of SR-Dyna and Dyna-Q. Median value function (grayscale) and implied policy after each algorithm (row) learns about relevant change in each of the 3 tasks (column). Both SR-Dyna (a) and Dyna-Q (b) can solve all 3 tasks when a sufficient number of samples backed up. c) Without a sufficient number of samples, SR-Dyna can still solve the latent learning task. d) Without a sufficient number of samples, Dyna-Q cannot solve any of the 3 tasks.

### Time constraints can distinguish SR-Dyna from Dyna-Q

SR-Dyna is capable of behaving equivalently to dynamic programming and tree-search in that it can solve transition and policy revaluation tasks. We were thus interested in how it could be differentiated experimentally. Sutton’s original Dyna algorithm (Dyna-Q) differs from value iteration and tree search in that its ability to pass revaluation experiments is dependent on having enough time offline to perform sufficient number of sample backups. Given enough time between decisions, sufficient replay can occur and Dyna-Q can pass any type of revaluation task (Fig 6b). In contrast, without sufficient offline replay, Dyna-Q degrades to a model-free agent and it cannot pass any revaluation task (Fig 6d). SR-Dyna is similarly differentiated from tree-search and value iteration in that its flexibility depends on a completing a sufficient number of sample backups offline. As demonstrated above, given a sufficient number of backups, SR-Dyna can pass any type of revaluation (Fig 6a). Without sufficient replay, its performance degrades to that of SR-TD – it can pass reward revaluation but fails transition and policy revaluation (Fig 6c). SR-Dyna is thus differentiated from Dyna-Q in that, unlike Dyna-Q, without sufficient replay, it can still pass reward revaluation; that is, it retains a certain degree of “model-based” flexibility even in the degraded case. These predictions could be tested with experimental designs aimed at preventing replay by manipulating the length of rest periods or the difficulty of distractor tasks [27].

Time constraints or distractor tasks at decision time can also disambiguate the different algorithms. Tree-search and value iteration take time and effort at decision time, whereas SR-Dyna can support rapid action selection by inspecting its lookup table.

## Interim Discussion

### Biological plausibility

Following real experience, SR-Dyna uses a similar update rule as SR-TD, yet uses it to operate over state-actions rather than states. This is plausible given that the same Hebbian learning principles could operate over cortical or hippocampal representations of state/action conjunctions just as well as they could over states.

As with Dyna-Q, SR-Dyna dovetails nicely with neural evidence about memory replay. Specifically, the widely demonstrated phenomenon of reactivation during rest or sleep of sequences of hippocampal activity seen during prior experiences [61,62], seems well suited to support the sort of off-line updates imagined by both Dyna approaches. (Although we have not simulated realistic hippocampal replay dynamics here, the Dyna approaches can learn from experiences replayed in arbitrary order.) The successor matrix updated by SR-Dyna might itself exist in the recurrent connections of hippocampal neurons [18], though another intriguing possibility is that it is instead stored in prefrontal cortex (as in Figure 1b). This second possibility lines up neatly with complementary system theories in the memory literature, according to which such hippocampal replay plays a role in post-encoding consolidation of memories by restructuring how information is represented across neocortical networks [63,64]. Such a connection should be explored in future research.

### Behavioral adequacy

Given sufficient replay, SR-Dyna is capable of producing behavior as flexible as that of full model-based value iteration. As with Dyna-Q, practical applications of SR-Dyna in larger environments will require developing sophisticated methods for selecting which samples to replay (e.g. [65]). We intend to develop such methods in future work.

## Discussion

Despite evidence that animals engage in flexible behaviors suggestive of model-based planning, we have little knowledge of how these computations are actually performed in the brain. Indeed, what evidence we have – particularly concerning the involvement of dopamine in these computations – seems difficult to reconcile with the standard abstract computational picture of planning by tree search using a learned model. We have here proposed variants of the SR that can address this question, serving as empirically consistent mechanisms for some or indeed all of the behaviors associated with model-based learning. Moreover, these are each built as utilizing a common TD learning stage for reward expectancies, allowing them to fit within the systems-level picture suggested by rodent lesion studies, and also explaining the involvement of dopamine in model-based valuation. In particular, they each envision how model-based learning could arise from the same dopaminergic TD learning associated with simple model-free learning, operating over a different and more elaborated cortical input representation.

### Accounting for TD circuitry in apparently model-based behavior

More specifically, our motivation to develop this approach was based on three related sets of findings in the empirical literature. The first are that lesions to dorsomedial striatum prevent animals from adjusting preferences following reward revaluation [5]. In contrast, lesions to neighboring dorsolateral striatum cause rats to maintain devaluation sensitivity, even following overtraining [38]. In the framework presented here, neurons in dorsomedial striatum could represent values derived by applying TD learning to the successor representation and neurons in dorsolateral striatum could represent values derived by applying TD to tabular representations. Lesions to dorsomedial striatum would thus force the animal to work with values in dorsolateral striatum, derived from tabular representations and thus not sensitive to devaluation. In contrast, lesions to dorsolateral striatum would cause the brain to work with values derived from the SR, which are devaluation-sensitive.

The second set of findings include several reports that the phasic DA response (or analogous prediction error related BOLD signals in humans) tracks apparently model-based information [6,11]. We have focused our simulations on choice behavior, and have not presented our theories’ analogous predictions about the responses of neurons, such as DA cells, thought to signal decision variables. However, whenever the SR algorithms’ expectations about action values incorporate “model-based” information (such as latent learning, Fig 4a) neural signals related to those predictions and to prediction errors would be similarly informed. Thus the theories predict systematic expectancy-related effects in the modeled dopamine response, tracking the differences in choice preference relative to the standard “model-free” accounts, which are blind to reward contingencies in these tasks.

A third distinct set of findings also speaks to a relationship between dopamine and model-based learning. These are reports that several measures of dopaminergic efficiency (both causal and correlational) track the degree to which human subjects engage in model-based decision strategies in both multistep reward revaluation tasks and multiplayer games [7–10,66]. One possibility is that these effects reflect strengthened vs. weakened phasic dopaminergic signaling, which in our model controls reward learning for SR-based “model-based” estimates in dorsomedial striatum. However, this account does not explain the specificity of these effects to measures of putative model-based (vs. model-free) learning. These effects may instead be related to functions of dopamine other than prediction error signaling, such as *tonic* dopamine’s involvement supporting working memory [67] or its hypothesized role controlling the allocation of cognitive effort [39,68,69].

### Other potential explanations

The framework outlined in this paper is not the only direction toward a neurobiologically explicit theory of putatively model-based behavior, nor even the only suggestion explaining the involvement of dopamine. As discussed above and pointed out in [27], Sutton’s original Dyna algorithm – in which experience replayed offline is used to update action values *V* or *Q* directly – offers another avenue by which seemingly model-based flexibility can be built on the foundation of the standard prediction error model of dopamine. This is a promising piece of the puzzle, but exclusive reliance on replay to underpin all behavioral flexibility seems unrealistic. Among our innovations here is to suggest that replay can also be used to learn and update a successor representation, which then confers many of the other advantages of model-based learning (such as flexibility in the face of reward devaluation) without the dependence on further replay to replan. Furthermore, the addition of SR to the Dyna framework explains a number of phenomena that replay, on its own, does not. For instance, given that Dyna-Q works with a single set of cached Q values, updated through both experience and replay, it is not clear how it could, on its own, explain the apparent segregation of revaluation sensitive and insensitive value estimates in dorsomedial and dorsolateral striatum [5,38].

Another potential solution to some of the puzzles motivating this work is that dopamine could have a role in action selection, as part of a circuit for partially model-based action evaluation [47]. According to this idea, dopamine neurons could compute a prediction error measuring the difference between the value of the current state and the future value of a predicted successor state, caused by a given candidate action. The size of this prediction error could then determine whether the action is performed. This mechanism would endow the brain with a single step of model-based prediction. However, it is not straightforward how this sort of approach could underlie model-based learning in tasks requiring more than a single step of prediction, and accordingly our simulations (see Figure 4a and supplemental materials) show that it cannot solve any of the revaluation tasks considered here, which all probe for deeper search through the state space. A recent study provided convincing behavioral evidence that humans sometimes simplify model-based action selection by combining just one step of state prediction with cached successor state values [70]. Yet this same study along with others [50] have also provided evidence that humans can plan through more than one step and thus are not confined to this approximation. It is also not straightforward how this sort of mechanism could endow model-based predictions in cases where stochasticity requires consideration of “trees” of possible future states.

Nevertheless, by elucidating a more general framework in which a predictive state representation may feed into downstream dopaminergic reward learning, we view our framework as fleshing out the spirit of this suggestion while also addressing these issues. We similarly realize other conceptual suggestions in the literature suggesting that more flexible model-based like behavior may arise not through tree-search like planning, but rather by applying model-free RL to more sophisticated state representations [71]. In a more specific application of this idea, [72] demonstrated that a sophisticated representation that includes reward history can produce model-based like behavior in the two-step reward revaluation task. The successor representation adds to this work by clarifying for any task’s transition structure, the precise representation that can be used to generate model-based behavior.

Relatedly, because it places model-based state prediction in the input to a standard TD learning circuit, our framework could easily be extended to include several modules with inputs corresponding to several different types or granularities of models: for instance, varying degrees of temporal abstraction corresponding to different time discount factors in equation 6 [44,73]. This would parallel a number of other recent suggestions that different striatal loops model the world at different levels of hierarchical abstraction [46,74], while also harmonizing the somewhat underspecified model-based evaluation process these theories assume with the predominant temporal difference account of striatal learning.

### Multiplicity and Arbitration

Although our presentation culminated with proposing an algorithm (SR-Dyna) that can in principle perform equivalently to full model-based learning using value iteration, this need not be the only goal and there need not be only a single answer. The behaviors associated with model-based learning may not have unitary sources in the brain but may instead be multiply determined. All of the algorithms we have considered are viable candidate pieces of a larger set of decision systems. Notably, the experiments we have highlighted as suggesting striatal or dopaminergic involvement in “model-based” learning and inspiring the present work all use extremely shallow planning problems (e.g. operant lever pressing, two-stimulus Pavlovian sequences, or two-step MDPs) together with reward revaluation designs. Even SR-TD is sufficient to explain these. It may well be that planning in other tasks, like chess, or in spatial mazes, is supported by entirely different circuits that really do implement something like tree search; or that they differentially require replay, like SR-Dyna. Also, although replay-based approaches go a long way, value computation at choice time using more traditional model-based approaches is likely needed at the very least to explain the ability to evaluate truly novel options (like the value of “tea jelly”; [75]) using semantic knowledge. Some evidence that rodents may use more than just replay to compute values, even in spatial tasks, comes from findings that the prevalence of sharp-wave-ripples, a putative sign of replay, is inversely related to the prevalence of vicarious trial and error behaviors, a process thought to be involved in decision-time value computation, potentially by standard MB dynamic programming or alternatively SR-MB [76].

Relatedly, if the brain might cache both endpoint decision variables like *Q*, or their precursors like *M*, update either or both with off-line replay, and optionally engage in further model-based recomputation at choice time, then the arbitration or control question of how the brain prioritizes all this computation to harvest rewards most effectively and efficiently becomes substantially more complicated than previously considered. The prioritization of replay – which memories to replay when – becomes particularly important. The particular ordering and dynamics of replay are also outside our modeling here: in order to focus our investigation on the simplest behavioral predictions of SR-Dyna, we chose the simplest, naive sampling scheme in which the agent replays a single state-action-state transition with uniformly random probability. This sampling strategy is not a mechanistic commitment (nor of course does it reflect the dynamics of realistic hippocampal replay trajectories), and we expect that like Dyna-Q, SR-Dyna would work even more efficiently given more sophisticated replay prioritization schemes. In this regard, we expect that the prioritization of replay, like the arbitration between model-based vs model-free tradeoffs [1,39,77], might operate according to the principles of efficient cost-benefit management. We expect that in addition to more typical observed patterns of replay, such a scheme may be able to explain cases where the replayed sequences are not a simple reflection of the animal’s current policy [78]. The current model is robust to differences in replay but would need to be extended with a more principled and detailed replay model to address these questions.

### Future Experimental Work

With simulations, we have presented experiments that could be used to elicit recognizable behavior form the different algorithms proposed here. Although we ruled out the simplest approach, SR-TD, due to its inflexibility, it is worth more carefully considering the evidence against it. The main counterexamples to SR-TD are transition revaluation and detour tasks. Apart from the classic work of Tolman and Honzik [55], the original results of which are actually quite mixed (see [79]), there is surprisingly little evidence to go on. A number of different studies have shown that healthy animals will normally choose the shortest alternative route after learning about a blockade preventing a previously preferred route (e.g. [80–82]). However, in these studies, the animal learns about the blockade after starting from the maze starting location. Thus, unlike in our simulations in which the animal learns about the blockade in isolation, animals in these tasks would have the opportunity to learn from direct experience that maze locations leading up to the blockade are no longer followed by maze locations further along the previously preferred path. Such tasks could thus potentially be solved by SR-TD. Studies that show that animals will take a shortcut to a goal that is discovered along a preferred path present a somewhat cleaner test for SR-TD [83,84]; however it is often difficult to interpret a potential role of exploration or visual (rather than cognitive map) guidance in the resulting behavior. Work in humans, however, seems to more clearly suggest an ability to solve detour tasks without re-learning [85]. Simon and Daw [23] for instance directly assessed SR-TD’s fit to human subjects’ choice adjustments in a changing spatial maze, and found it fit poorly relative to traditional model-based learning.

Overall, additional careful work that measures how animals respond to transition changes, learned in isolation, is needed. Whereas Tolman’s other early reward revaluation experiments (latent learning) have been conceptually replicated in many modern, non-spatial tasks like instrumental reward devaluation and sensory preconditioning, the same is not true of detours. Indeed, the modern operant task that is often presented as analogous to detours, so-called instrumental contingency degradation (e.g., [86]), is not functionally equivalent. In such tasks, the association between an action and its outcome is degraded through introduction of background rewards. However, because the information about the changed contingency is not presented separately from the rest of the experience about actions and their rewards, unlike all the other tests discussed here, contingency degradation as it has been studied in instrumental conditioning can actually be solved by a simple model-free learner that re-learns the new action values. The puzzle here is actually not how animals can solve the task, but why they should ever fail to solve it. This has thus led to a critique not of model-based but of model-free learning theories [46].

In any case, the modeling considerations proposed here suggest that more careful laboratory work on “transition revaluation” type changes to detect use of SR-TD, is warranted. Similarly, “policy revaluations” along the lines of that in Fig 5 would be useful to detect to what extent planning along the lines of SR-MB is contributing. Finally, although SR-Dyna in principle can perform model-based value computation, this depends on sufficient replay. The SR-Dyna hypothesis suggests the testable prediction that behavior should degrade to SR-TD under conditions when replay can contribute less. A number of experiments in the rodent literature have explored the behavioral deficits that result from interrupting sharp-wave ripples (events in which hippocampal replay is known to occur). Such manipulations have been shown to produce behavioral deficits that are consistent with the SR-Dyna hypothesis, yet not exclusively predicted by it. For instance, two studies found that suppression of hippocampal sharp-wave ripples during rest slows down acquisition of the correct behavioral policy in spatial learning tasks in which the task environment is static [87,88]. These results are consistent with the notion that the purpose of replay is to provide additional experience, which is used to update some representation relevant to learning. However, these results are not specifically diagnostic of the successor matrix. For instance, preventing replay would slow down policy acquisition for Dyna-Q as well as SR-Dyna (Supplementary Figure 2).

Another study found a more specific effect of suppressing hippocampal sharp wave ripples during performance of a task. Here, the manipulation caused learning deficits selective for a subset of trials in which animals faced a hidden-state problem. In particular, these were trials in which animals had to choose which direction to turn depending on the events of the previous trial [89]. Such tasks constitute hidden-state problems, in that the “state” required to make the correct choice cannot be deduced entirely from an animal’s immediate sensory experience. In RL terms, to solve these problems, the animals must construct an augmented internal state, distinct from a simple representation of the immediate sensory situation [90]. One interpretation of the experimental result is that blocking replay interfered with this internal state construction process.

This result resonates with our SR-Dyna proposal, which also posits that replay is involved in constructing the mapping between the sensory state and a different, augmented internal representation of it: the SR. However, our model as currently specified augments the state space with predictive features to support model-based flexibility, and does not currently address other sorts of elaborations of the state input that have been used in other work to facilitate learning in situations with hidden state or other uncertain sensory input [91–93]. Fully understanding these results therefore requires augmenting our model to address hidden state as well as state prediction. In fact, these two functions may be closely related: a number of approaches to the hidden state problem in the computational RL literature address it using predictive representations that are related to the SR [94,95].

In human studies, factors like the duration of off-task rest periods and presence of distractor tasks during such periods have been manipulated to extend, limit, or interfere with replay. Some evidence suggests that distractor tasks at decision time have no effect on reward revaluation [27], consistent with SR-Dyna. Other recent work has demonstrated that humans benefit from additional pre-decision time in revaluation tasks that closely resemble “policy revaluation” [96] and that this benefit recruits a network including the prefrontal cortex and basal ganglia. Such work is consistent with the predictions of both Dyna-Q as well as SR-Dyna accounts of value updating presented here.

Overall, future work will need to combine such manipulations of replay, with the three revaluation tasks imagined in this paper and demonstrate differences in the effects of manipulations on reward versus transition and policy revaluations. We have recently demonstrated, though without attempting to manipulate replay, that humans are worse at adjusting behavior following transition and policy revaluations compared to reward revaluations, suggesting that they may at least partially use either an SR-TD or SR-Dyna (with limited sample backups) strategy for evaluation [57].

### Neural substrate of the successor representation

We have suggested, on the basis of rodent lesion studies, that the SR may be encoded in parts of prefrontal cortex that project to dorosomedial striatum. However, we should note that recent work has also implicated the hippocampus as a potential site of the SR. Specifically, a state-state version of the SR can explain some properties of hippocampal place cells [18,97] as well as fMRI measures of the representation of visual stimuli in tasks where such stimuli are presented sequentially [98,99]. This work has largely built on ideas of the hippocampus in general as a site of cognitive map [100] as well as prior suggestions that hippocampal place cells may in fact encode the transition structure of the environment [101] and that such transition information may make them ideal basis functions for TD learning [102] If a state version of the SR exists in the hippocampus, we think it is reasonable that value weights would be leaned by neurons connecting the hippocampus to ventral striatum, in the same TD manner discussed in this paper.

However, we also think a case can be made for the prefrontal cortex as another candidate basis for an SR. (These proposals are in no way mutually exclusive.) In addition to the rodent lesion evidence reviewed in the introduction of this paper, the prefrontal cortex shares many cognitivemap properties observed in the hippocampus [103] and has been suggested to be the basis of state representations for reinforcement learning [91,104]. A number of human studies have demonstrated the PFC’s role in the representation of prospective goals [105,106]. Furthermore, unlike the hippocampus, parts of the prefrontal cortex appear to be involved in action representation in addition to state representation [107], thus making it a candidate to hold a potential state-action version of the successor matrix. Overall, further experimental work will be required to determine whether either or indeed both these areas serves as the basis for the successor representation, and what specific roles they play in learning and representation.

### Connection to other cognitive processes

Finally, the SR may contribute to a number of other cognitive processes. Above we noted that there is evidence that areas of medial temporal lobe seem to encode predictive representations. In line with this, it has been noted that there is a close correspondence between the update rule used by SR-TD and update rules in the temporal context model of memory [19]. Also, recent approaches to reinforcement learning in the brain have advocated for a hierarchical approach in which punctate actions are supplemented by temporally abstract policies [108]. In this context, it has been suggested that the SR may be useful for discovering useful temporal abstractions by identifying bottlenecks in the state space that can then be used to organize states and action into a hierarchy [18,109]. The efficacy of the SR for model-based RL opens the possibility that the brain accomplishes planning, action chunking, and grouping episodic memories using a common mechanism.

Overall, this article has laid out a family of candidate mechanistic hypotheses for explaining the full range of behaviors typically associated with model-based learning, while connecting them with the circuitry for model-free learning as currently understood. In addition to the transition and policy revaluation behavioral experiments suggested above, future neuroimaging work could seek evidence for these hypotheses. Specifically, failures to flexibly update decision policies that are caused by caching of either the successor representation (as in SR-TD or SR-Dyna with insufficient replay) or a decision policy (as in SR-MB) should be accompanied by neural markers of non-updated future state occupancy predictions. Such neural markers could be identified using representational similarity analysis (e.g. [110]), cross-stimulus suppression (e.g. [111]) or through use of category specific, decodable, visual stimuli (e.g. [112]). Similar work in experimental animals such as rodents (e.g. [113]) could use the full range of invasive tools to trace the inputs to dorsomedial vs. dorsolateral striatum, so as to examine the information represented there and how it changes following the various sorts of revaluation manipulations discussed here. As has been the case for model-free learning, the emergence of an increasingly clear and quantitative taxonomy of different candidate algorithms is likely to guide this work and help to elucidate the neural basis of model-based learning.

## Methods

### General Simulation Methods

All simulations were carried out in 10x10 (N = 100 states) grid-worlds in which the agent could move in any of the four cardinal directions, unless a wall blocked such a movement. States with rewards contained a single action. Upon selecting that action, the agent received the reward and was taken to a terminal state. Each task was simulated with each algorithm 500 times. For each simulation, we recorded the agent’s value function at certain points. For SR-Dyna, which worked with action values rather than state values, the state value function was computed as the max action value available in that state. Figures display the median value, for each state, over the 500 runs. To determine the implied policy for the median value function, we computed, for each state, which accessible successor state had the maximum median value.

### Specific Task Procedures

#### Latent learning task

The latent learning task was simulated in the grid-world environment shown in Fig. 2a. Starting from position S, the agent first took 25000 steps exploring the maze. After exploration, the reward in position R1 was raised to 10. To learn about the reward, the agent completed a single step, starting from position R1, 20 times. We then recorded the state value function.

#### Detour task

The detour task was simulated using the grid-world environment shown in Fig. 2b. Starting from position S, the agent first took 10000 steps exploring the maze. The reward in position R was then increased to 10. The agent then completed 5 trials, starting from position S that ended when the reward was reached. A wall was then added to in position B. To learn about the wall, the agent completed a single step, starting from the position immediately left of the wall, 40 times. We then recorded the state value function.

#### Novel revaluation task

The novel revaluation task was simulated using the environment in Fig. 5c. The agent first completed the entire latent learning task. After successfully reaching position R1 from position S, the agent then completed 20 trials. Each trial alternately started at S1 or S2 and ended when the agent reached position R1. We then set the reward in position R2 to 20. To learn about the new reward, the agent completed one step, starting from position R2, 20 times. We then recorded the state value function.

### Additional Details on Algorithms

#### One-step look-ahead

The one-step look-ahead model stored an estimate of state-value function *V*^*π*^ *s*. At the beginning of each simulation *V*^*π*^ was initilzled to **0**. Following each transition *V*^*π*^ was updated according to equation 4. Prior to each choice, Q-values for each action *a* in state *s* were then computed as *Q*^*π*^ (*s*, *a*) = *V*^*π*^(*s*’) where *s*’ is the state that deterministically follows action a in state s. Note that leaving *R*(*s*, *a*) out of this equation works because rewards are paired exclusively with actions in terminal states (and thus *R*(*s*, *a*) for non-terminal actions is 0).

#### Original successor representation (SR-TD)

SR-TD computed *V*^*π*^(*s*) using two structures: the successor matrix, *M*^*π*^(*s*, *s*’) and a weight vector, *w*(*s*). At the beginning of each simulation, *M*^*π*^ was initialized as an identity matrix; however, rows corresponding to terminal states were set to 0. The weight vector was initialized as *w* = **0**. Following each transition, *M* and *w* were updated using equations (8) and (9). In implementing the update in equation 8, each element of the feature vector, *M*(*s*,:), was scaled by *M*(*s*,:)* *M* (*s*,:) ^*T*^. This scaling permits the weight learning rate parameter to maintain a consistent interpretation as proportional step-size. Prior to each choice, *V* was computed using equation (7). Q-values for each action were computed the same as for the one-step look-ahead model.

#### Recomputation of the successor matrix (SR-MB)

This algorithm starts each task with a basic knowledge of the ‘physics’ of grid-world: which successor state, *s*’, would follow each action *sa* in a situation in which *sa* is available (e.g. not blocked by a wall). It also stores and updates, for each state *s*, *A*_*s*_, the set of actions currently available in state *s* as well as a policy *π*(*a*|*s*), which stores the probability of selecting action *a* in state *s* (as learned from the agent’s own previous choices). *A*_*s*_ was initialized to reflect all four cardinal actions being available in each state. Each row *π* were initialized as a uniform distribution over state-actions, *π*(*a*|*s*) = 0.25.

After performing action *a* in state *s* and transitioning to state *s*’, *A*_*s’*_ was updated to reflect which actions are available in state *s*’ and *π* is updated using a delta rule:

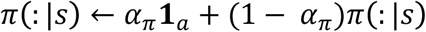

where *α*_*π*_ is a free parameter.

Prior to each choice, the model computed each row, s, of one-step transition matrix *T* ^*π*^ as follows:

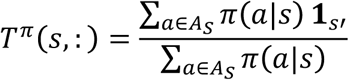

where **1**_*s*_ is a vector zeros of length *S* with a 1 in position corresponding to state *s*’ and *s*’ is the state to which action *a* in state *s* deterministically leads. *T* ^*π*^ was then used to compute *M* using equation 10. Computation of *V* and *Q* was then the same as in SR-TD.

#### Episodic replay algorithm (SR-Dyna)

This algorithm computed *Q*(*sa*) using two structures: a state-action successor matrix, *H*(*sa*, *s*’*a*’) and weight vector *w*(*sa*). At the beginning of each simulation, the successor matrix *H* was initialized to an identity matrix; however rows corresponding to terminal states were set to **0**. The weight vector was initialized to *w* = **0**. The algorithm also stored every sample (*s*, *a*, *s*’). After performing action *a* in state *s* and transitioning to state *s*’ the sample (*s*’, *a*’, *s*’) was stored, and *H* and *w* were updated according to equations (13) and (14). Following each step, we also selected 10 one-step samples (according to recency weighted probabilities with replacement) from the stored history, and replayed each to update H according to equation 15. Following transitions in which a learned change occurred to either the reward function or available actions, *k* one-step samples were selected and used to update the model, where k was set to 10 in the insufficient replay condition and to 10000 in the sufficient replay condition. Samples were drawn by first selecting a state-action, *sa*, from a uniform distribution. A sample then drawn from the set of experienced samples, initiated in *sa,* according to an in, initiated from sa was then selected according to an exponential distribution with *λ* = 1/5.

#### Dyna-Q

This algorithm stored *Q*(*sa*). At the beginning of each simulation, *Q* was initialized to **0.** The algorithm also stored every experienced sample (*s*, *a*, *r*, *s*’). After performing action *a* in state *s*, experiencing reward r, and transitioning to state *s*’ the sample (*s*’, *a*’, *r*, *s*’) was stored and Q was updated according to the Q-learning prediction error:

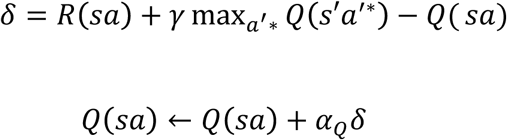

Following each step, as with SR-Dya, we also selected 10 one-step samples (according to recency weighted probabilities with replacement) from the stored history, and replayed each to update *Q* according to equation the update above. Following transitions in which a learned change occurred to either the reward function or available actions, *k* one-step samples were selected and used to update the model, where k was set to 10 in the insufficient replay condition and to 10000 in the sufficient replay condition. Samples were drawn the same way as in SR-Dyna.

#### Parameters

All algorithms converted Q-values to actions using an *ϵ*-greedy policy which selects the highest-valued action with probability 1 – *ϵ*, and chooses randomly with probability *ε*. Parameters for all models and each simulation were varied and we observed that the qualitative results can be observed under a wide range of parameter settings (Supplementary Table 1). For the figures in the results section, the following parameters were used. For all models, *ε* was set to 0.1. In addition, all models used a discount parameter *γ* = 0.95. The three SR models used a weight learning rate parameter *α*_*w*_ = 0.3. Model Dyna-Q used a learning rate *α*_*Q*_= 0.3. In addition to these parameters, SR-TD and SR-Dyna used a successor-matrix learning rate of *α*_*sr*_= 0.3 and SR-MB used a policy learning rate of *α*_*π*_= 0.1.

## Supplementary Materials

**Supplementary Figure 1:**
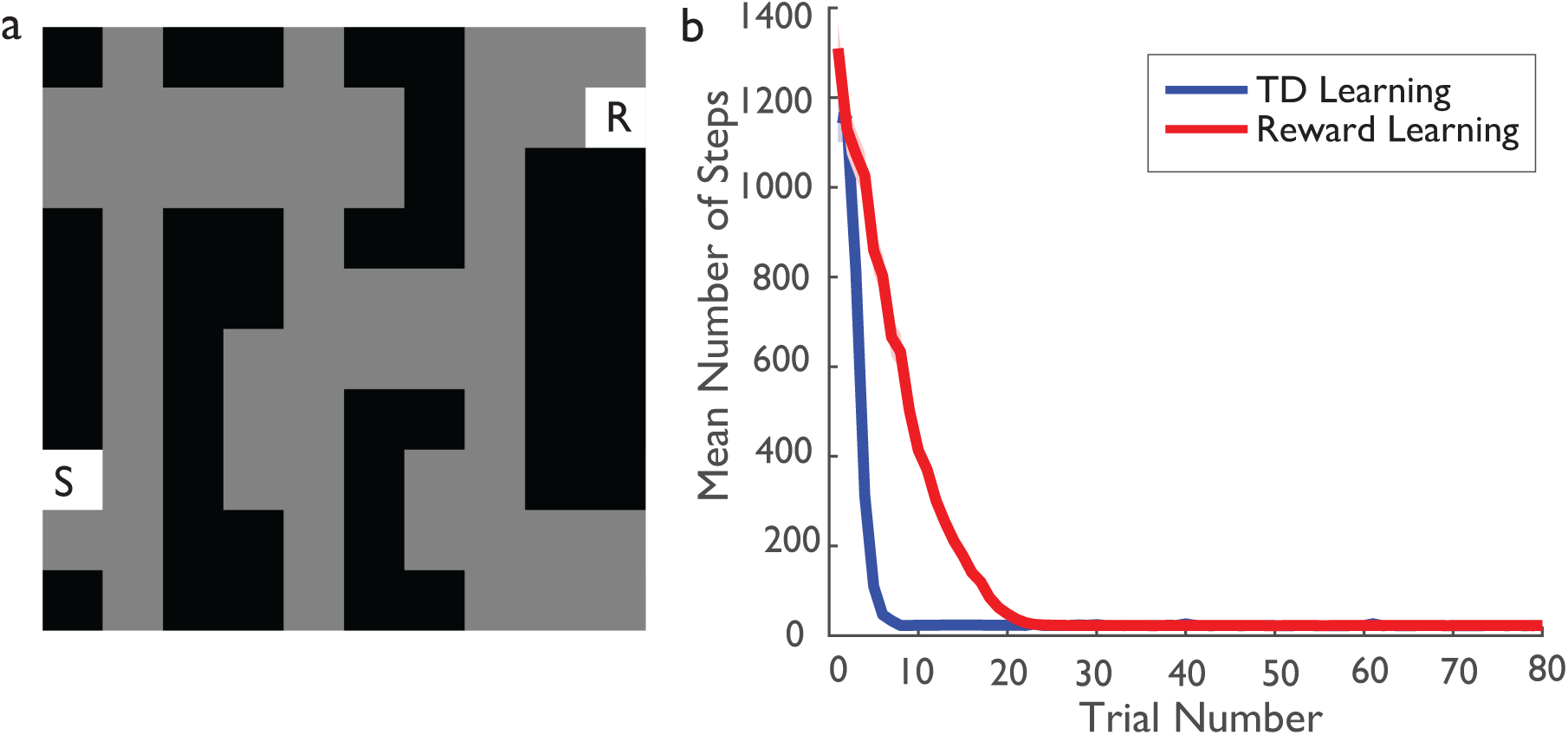
Advantage of TD learning over direct reward learning of weights. a) Task environment. On each trial, the agent was placed in state S. Trials ended when the agent reached state R, which contained a reward value of 10. Unlike the latent learning task in the main text, in this task did not contain an exploratory period enabling the agent to learn the successor matrix prior to introduction of reward. b) Number of steps on each trial for agent learning weights using TD learning and agent learning weights using reward learning. Plotted lines show average over 500 runs. 95% confidence intervals are contained within line thickness. Parameters for each of the two algorithms were set to those that minimized the average number of total steps over 80 trials. Such parameters were found by grid search over *α*_*sr*_ ∈ [. 1,.3,.5,.*7*,.9], *∈* ∈ [0.1, 0.3, 0.5] and *α*_*w*_ ∈.[1,.3,.5,.*7*,.9]. Both algorithms learned the SR using the SR-TD update. The “TD Learning” algorithm updated weights using TD learning. The “Reward Learning” algorithm updated weights by delta-rule learning on the immediate reward function. Specifically, after performing action *a* in state *s* and receiving reward *r*, the following update was performed: *w*(*s*)← *w*(*s*)+ *α*_*w*_(*r* – *w*(*s*)).

**Supplementary Figure 2:**
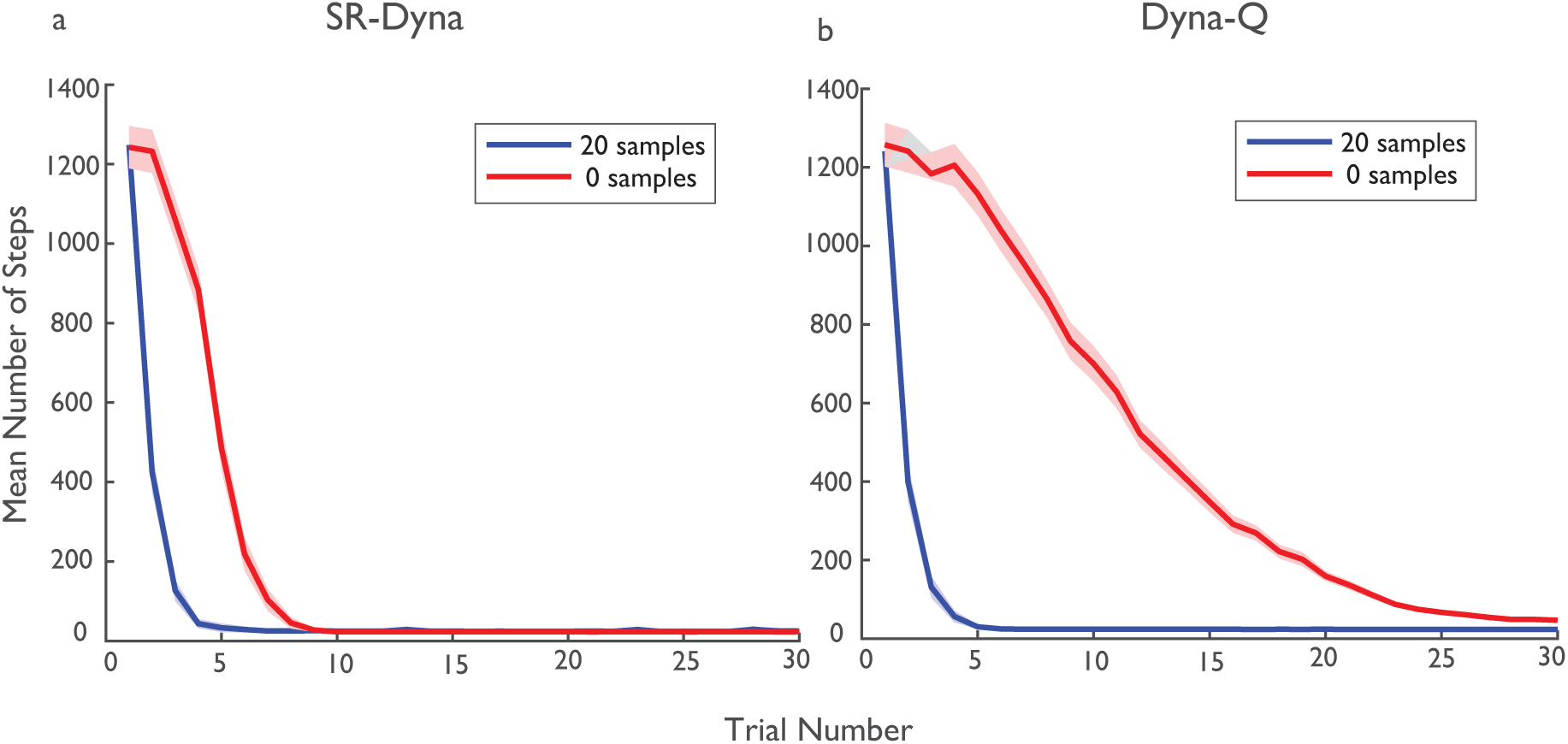
Preventing replay slows acquisition for both a) SR-Dyna and b) Dyna Q. Both algorithms under the two sampling settings were simulated on the task displayed in Supplementary Figure 1. Both a) and b) show number of steps on each trial for agent permitted to replay 20 samples between each decision and an agent not permitted to replay any samples. Plotted lines show average over 500 runs. 95% confidence intervals are contained within shaded region around lines. For each algorithm and sample setting, we chose parameters by a grid search in the following range: *α*_*sr*_ ∈ [. 1, .3, .5, .7, .9], ∈ ∈ [0.1, 0.3, 0.5], *α*_*w*_ ∈ [. 1, .3, .5, .7, .9], and *α*_*Q*_ ∈ [. 1, .3, .5, .7, .9].

**Table 1:**
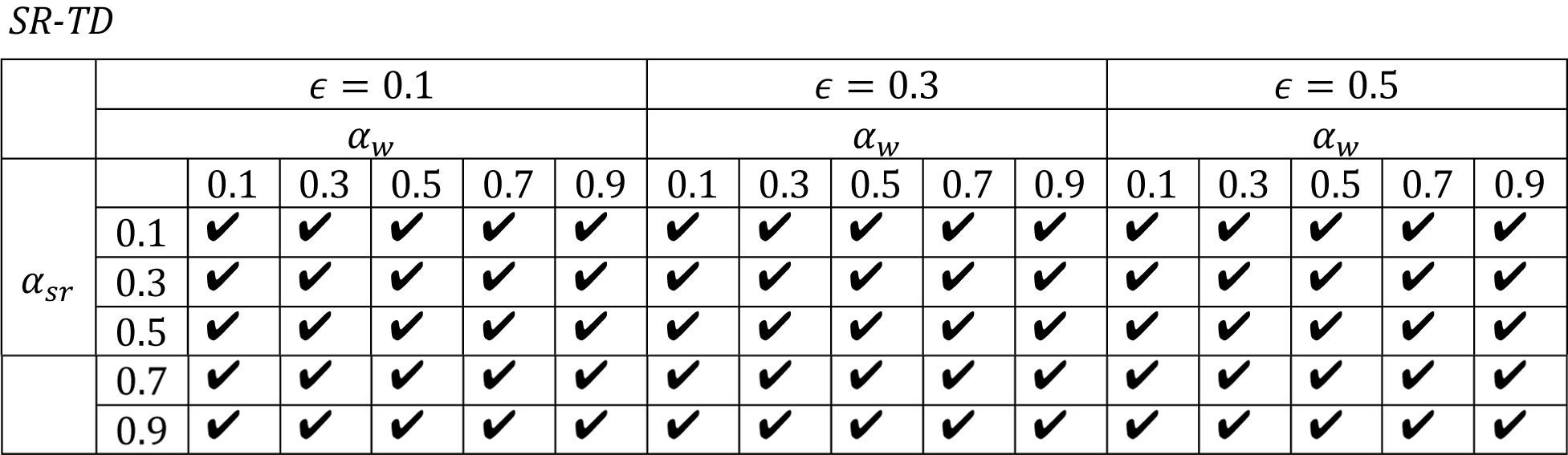
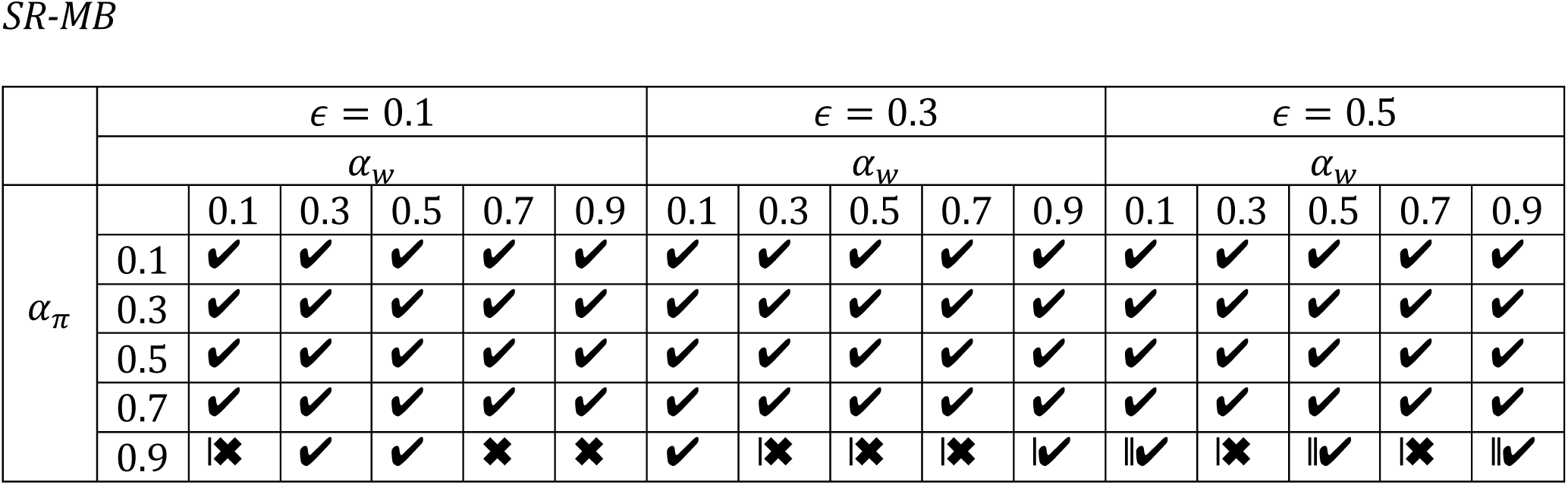
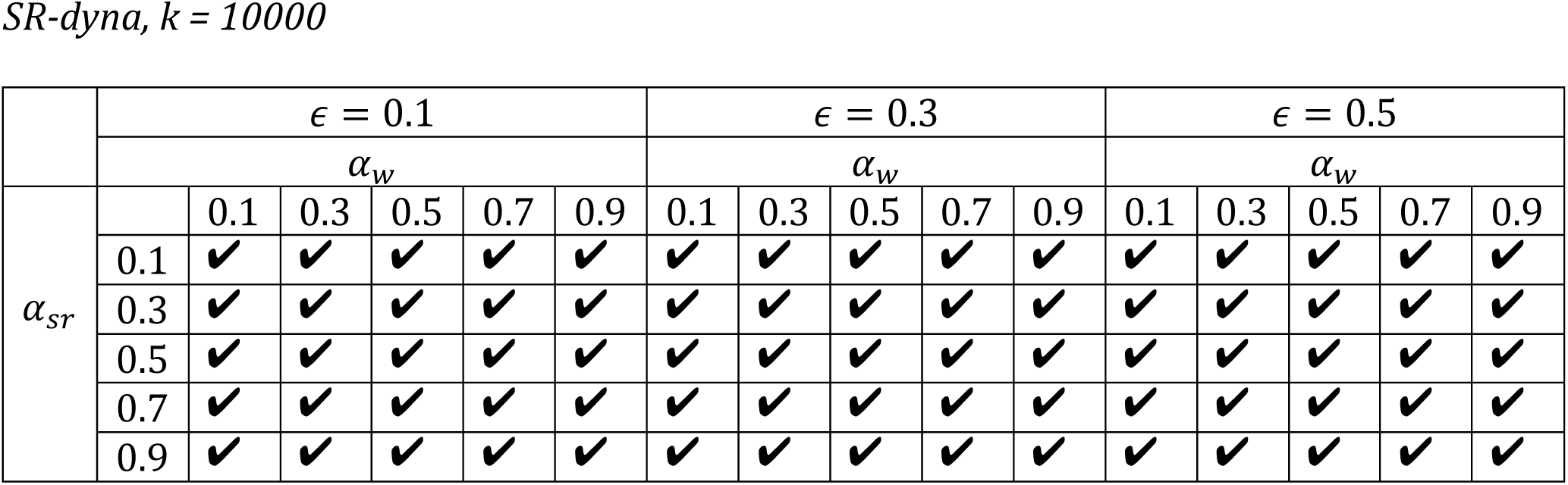
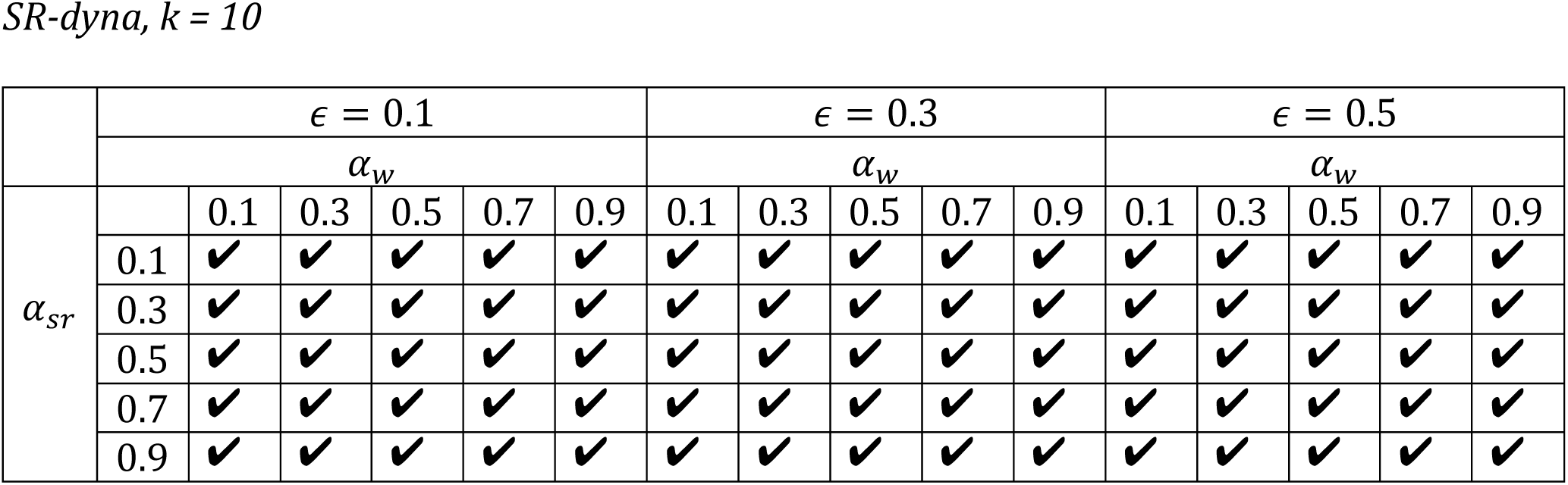
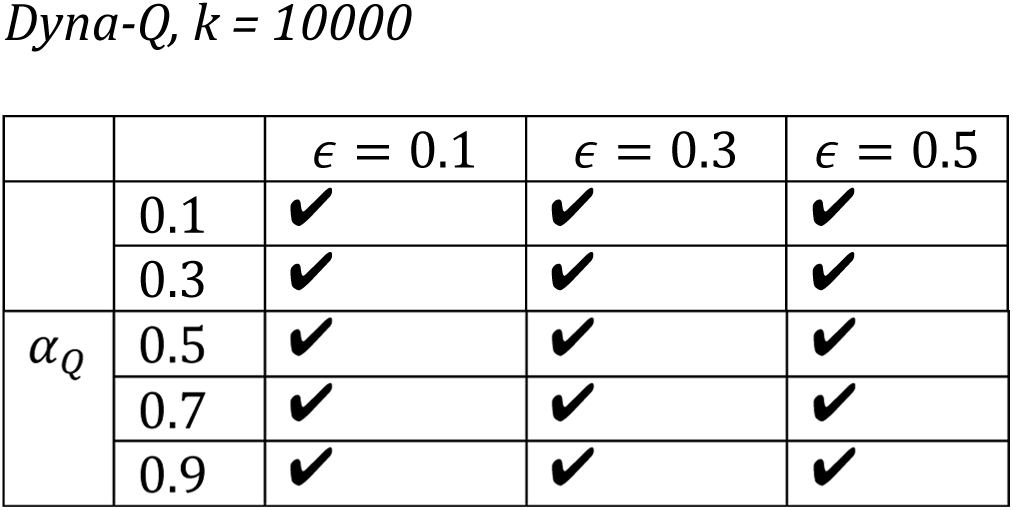
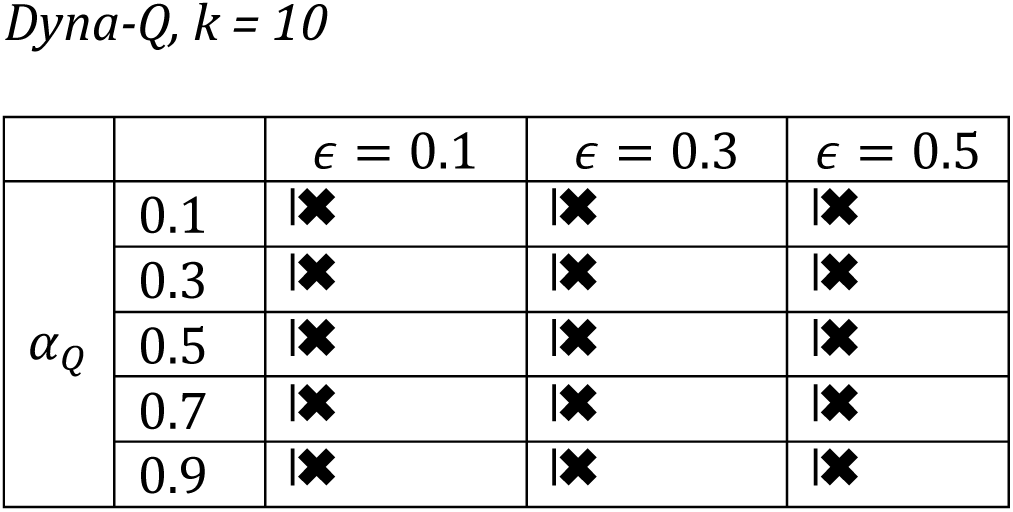
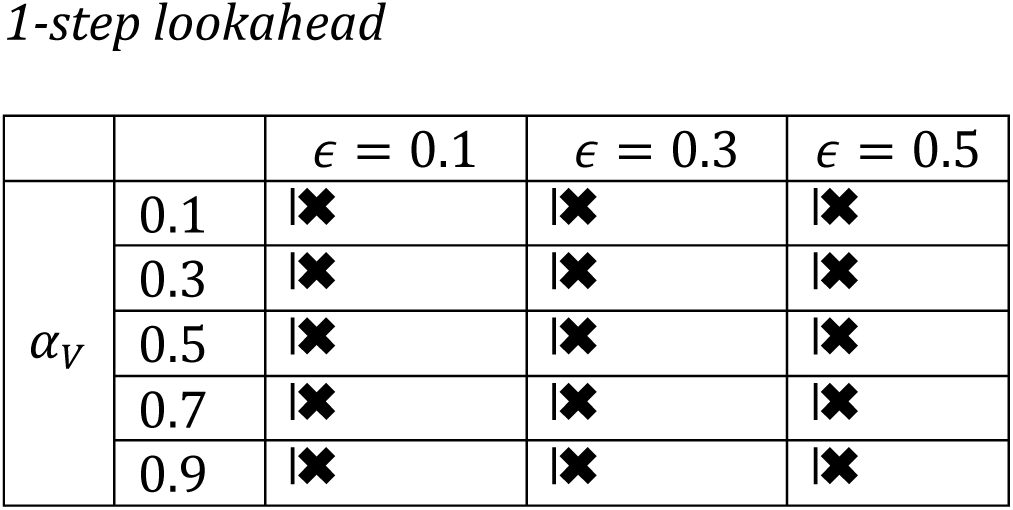
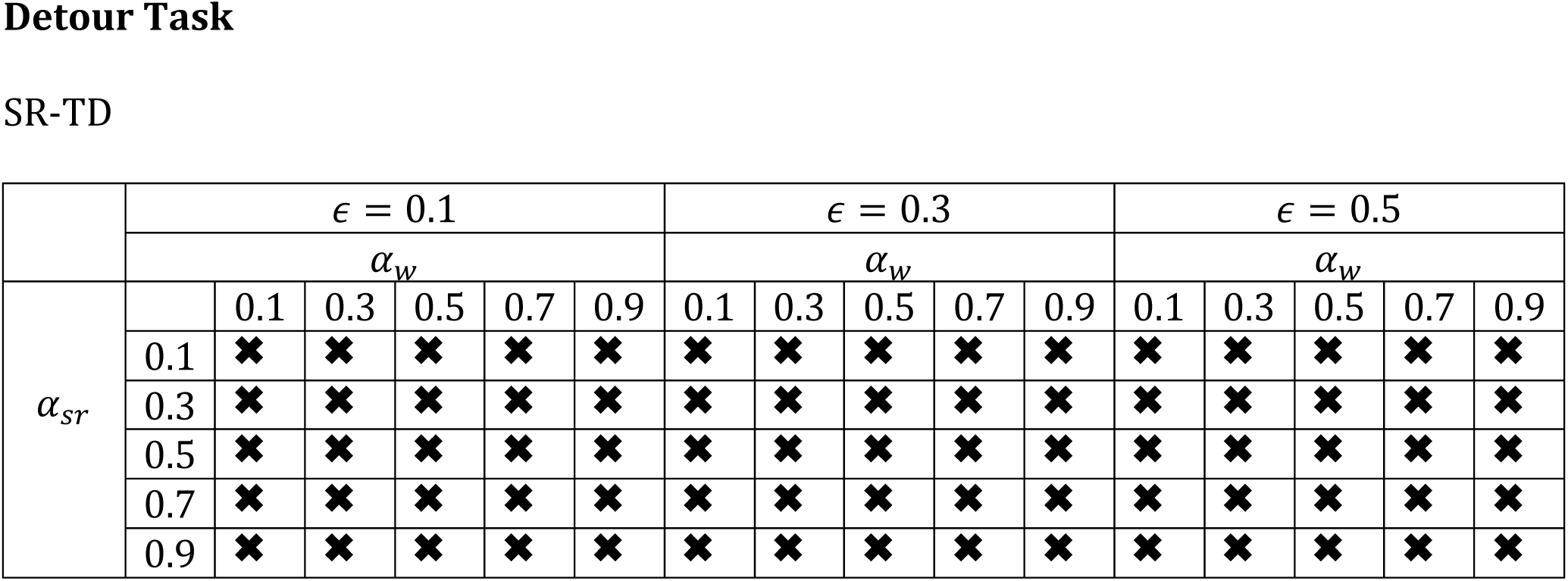
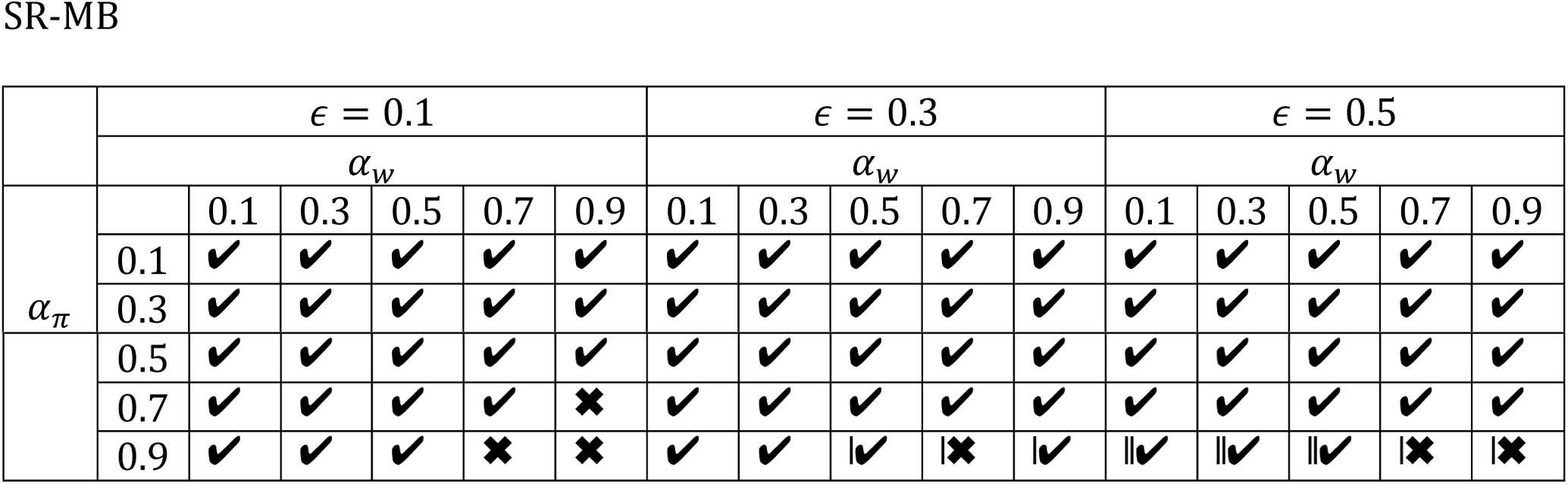
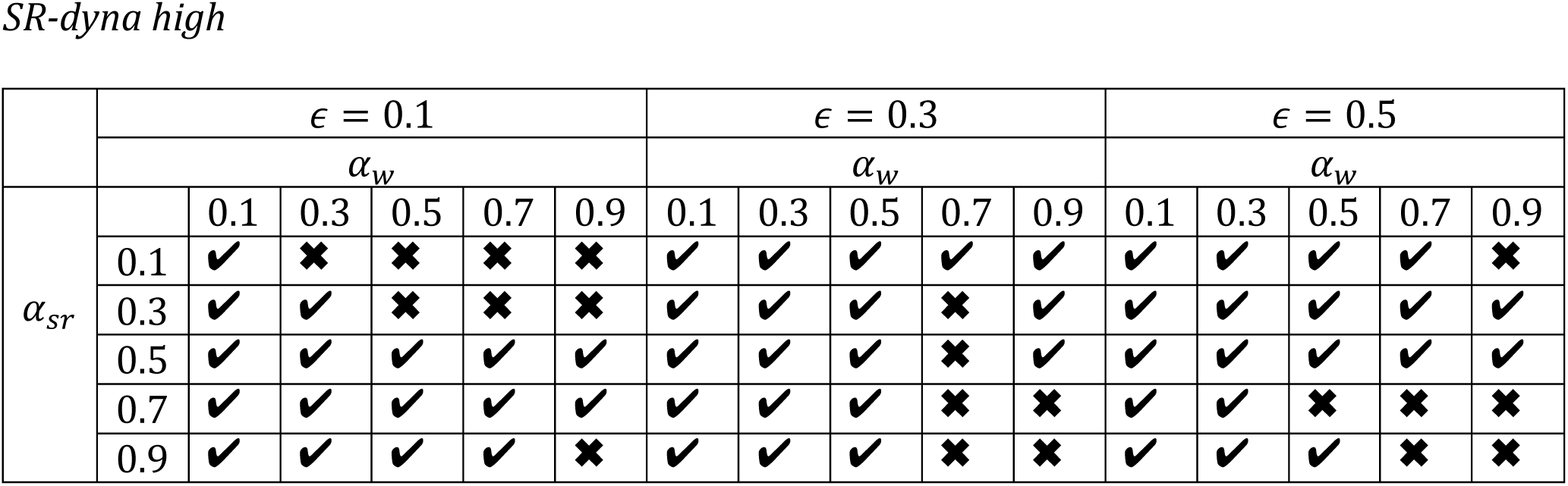
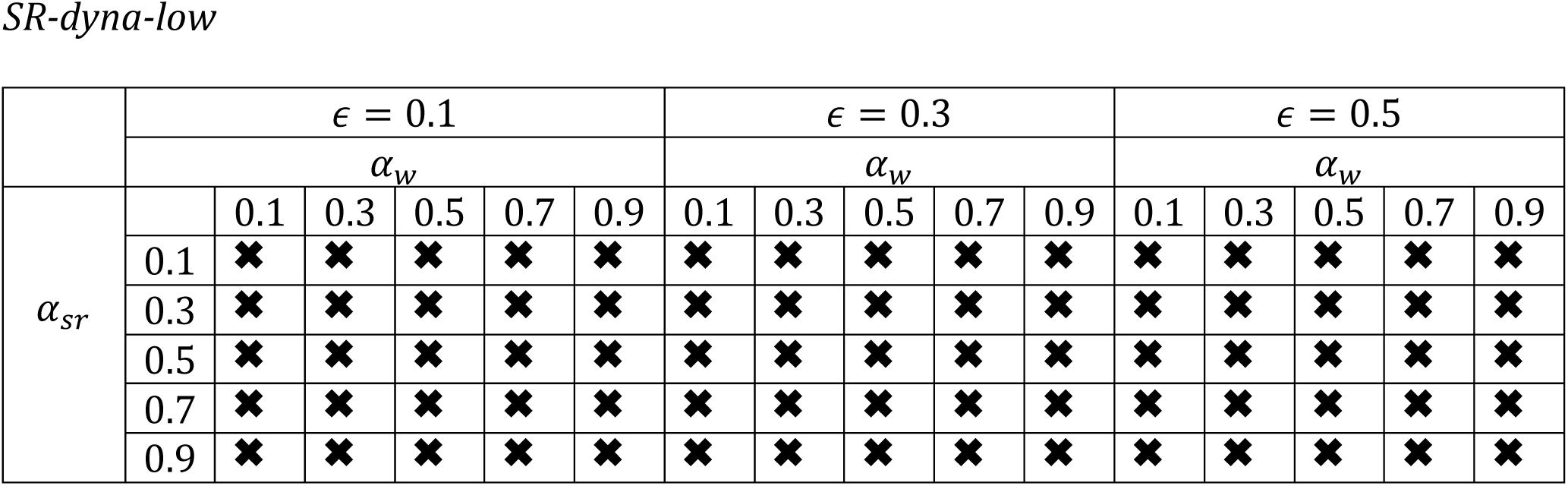
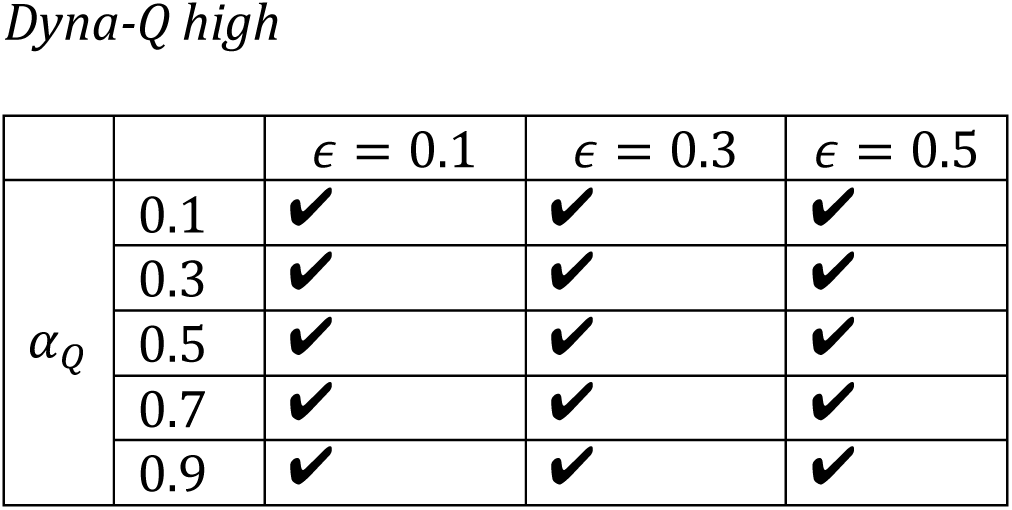
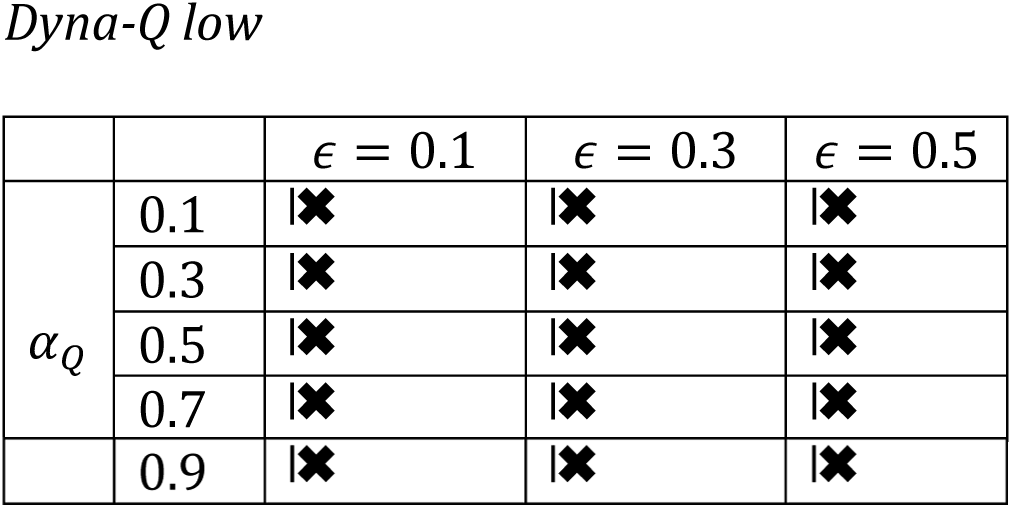
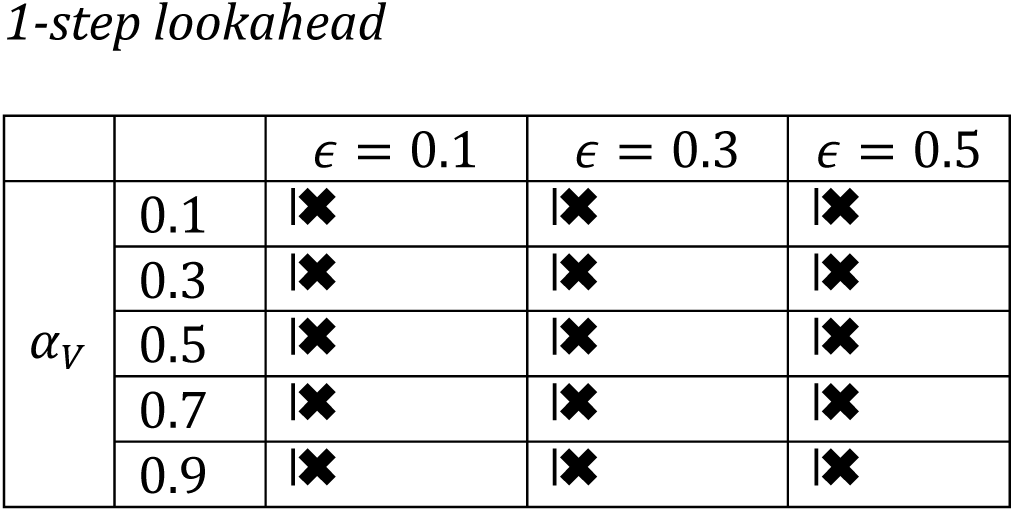
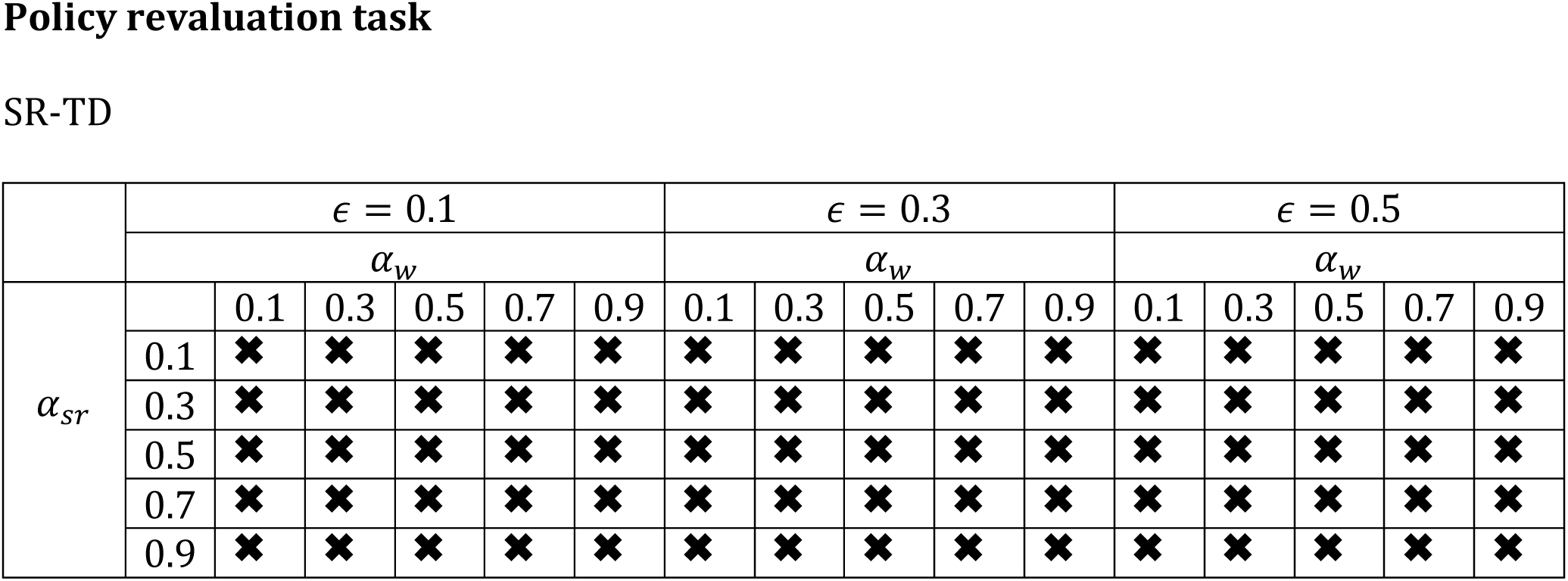
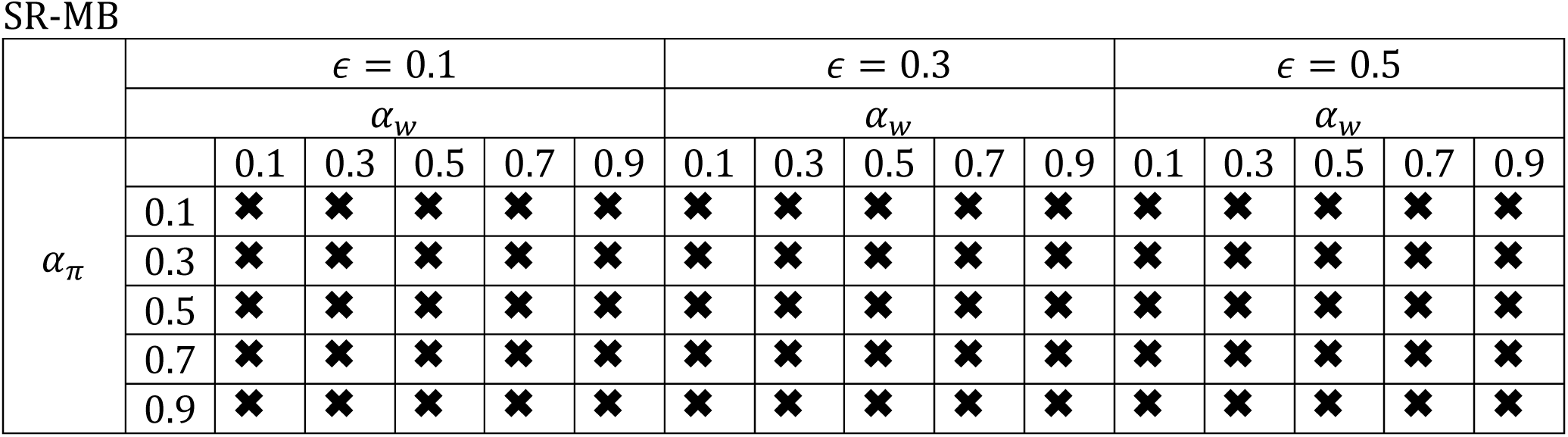
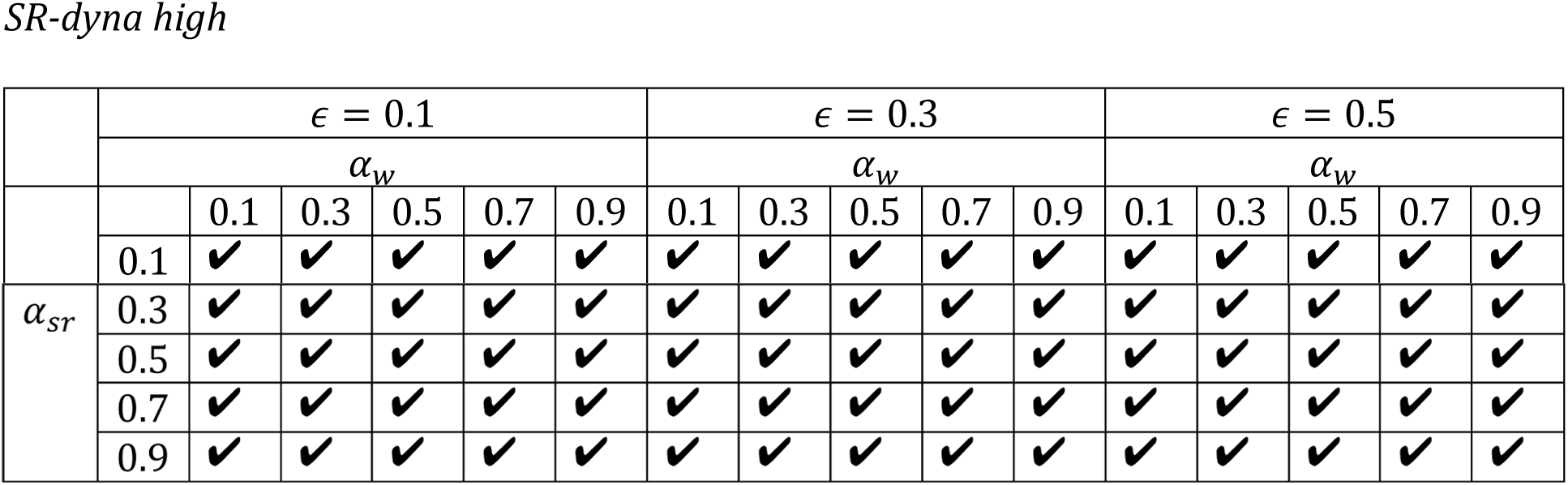
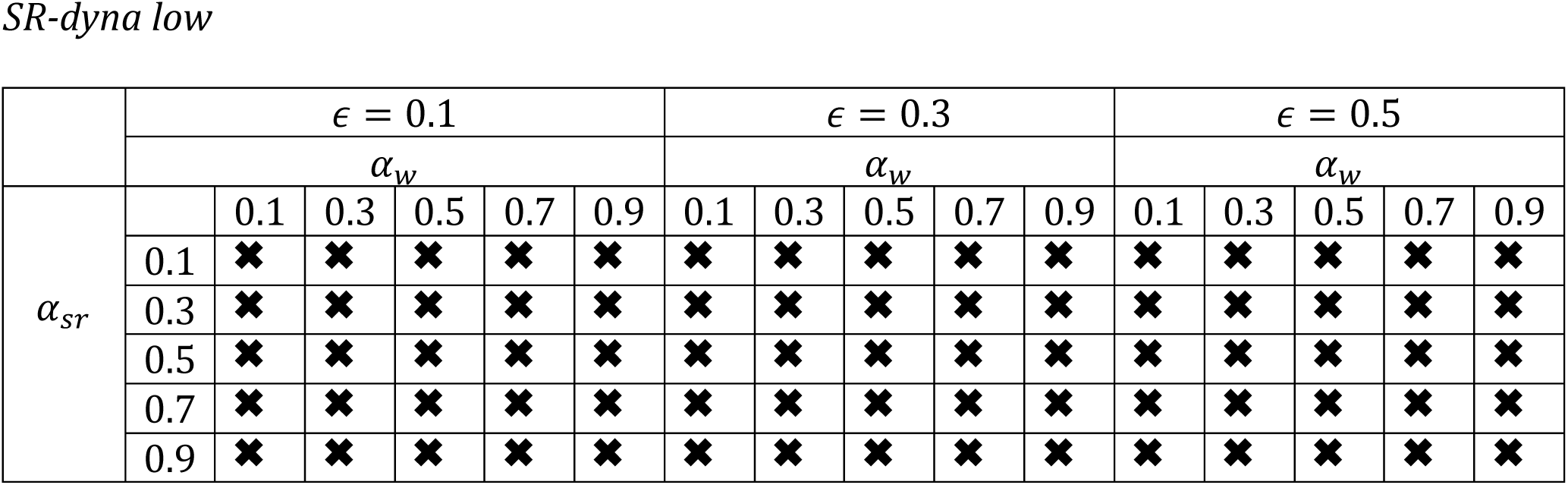
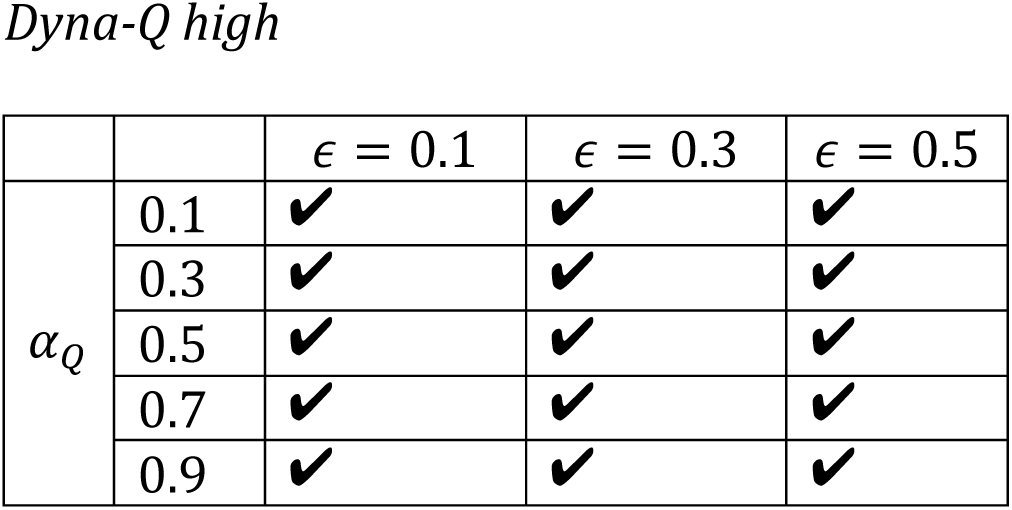
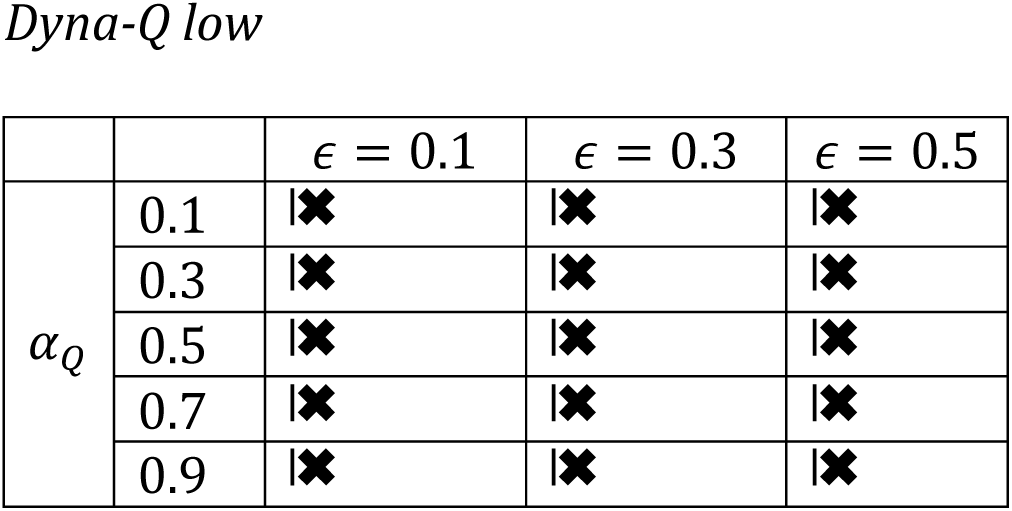
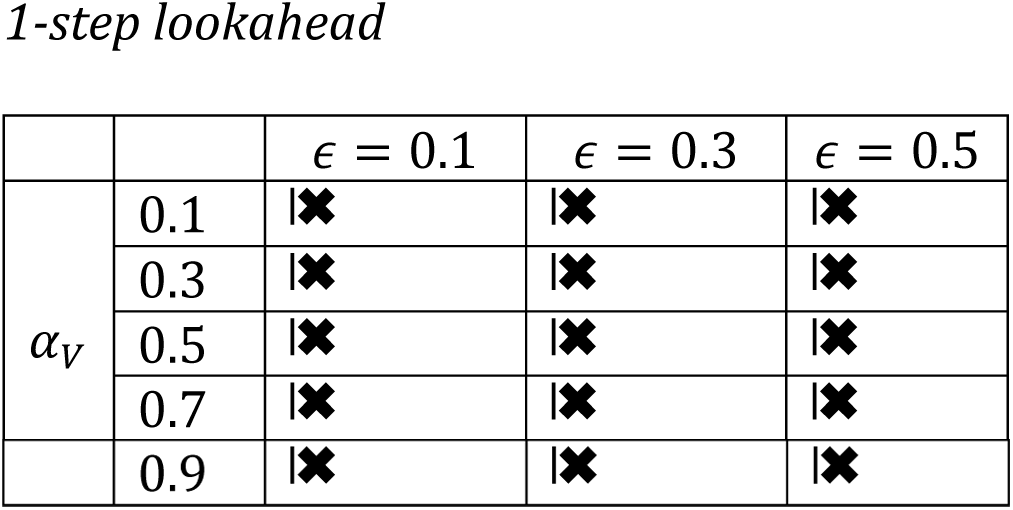
Robustness of simulation results to varying parameters. Here, we display the results of simulating each task, using each algorithm under a wide variety of parameter settings. Each table below corresponds to a particular algorithm simulating a particular task. For a given parameter setting, the algorithm was simulated 500 times. A check indicates that the 500 run median value function produced by that parameter setting results in the optimal policy for the task. A cross indicates that it does not result in the optimal policy. **Latent learning task**

